# State Space Methods for Phase Amplitude Coupling Analysis

**DOI:** 10.1101/772145

**Authors:** Hugo Soulat, Emily P. Stephen, Amanda M. Beck, Patrick L. Purdon

## Abstract

Phase Amplitude Coupling (PAC) is thought to play a fundamental role in the dynamic coordination of brain circuits and systems. There are however growing concerns that existing methods for PAC analysis are prone to error and misinterpretation. Improper frequency band selection can render true PAC undetectable, while non-linearities or abrupt changes in the signal can produce spurious PAC. Current methods require substantial amounts of data and lack formal statistical inference tools. We describe here a novel approach for PAC analysis that substantially addresses these problems. We use a state space model to estimate the component oscillations, avoiding problems with frequency band selection, nonlinearities, and sharp signal transitions. We represent cross-frequency coupling in parametric and time-varying forms to further improve statistical efficiency and estimate the posterior distribution of the coupling parameters to derive their credible intervals. We demonstrate the method using simulated data, rat LFP data, and human EEG data.

## 1 Introduction

Neural oscillations are thought to play a fundamental role in the dynamic coordination of brain circuits and systems [1]. At individual frequencies, oscillations reflect the temporal coordination of activity across populations of neurons, and can be observed experimentally in neuronal spiking time series, multi-unit activity, local field potentials (LFP), and even non-invasively using magnetoencephalogram or electroencephalogram (EEG) recordings. In the past decade, a major advance has been the realization that oscillating neural activity can have higher-order interactions in which oscillations at different frequencies interact [2–4]. This cross-frequency coupling (CFC) appears to be nearly as ubiquitous as oscillations themselves, occurring during learning and memory, varying across different states of arousal and unconsciousness, and changing in relation to neurological and psychiatric disorders [2,4–15]. If distinct oscillations stem from specific neural circuit architectures and time constants [16], it seems plausible that cross-frequency coupling could serve as a way of coordinating activity among otherwise disparate circuits and systems [2]. Amplitude-Amplitude [17] and Phase-Phase coupling [3] [18] have been reported, but phase-amplitude coupling (PAC), in which the phase of a slower wave modulates the amplitude of a faster one, remains the most frequently described phenomenon.

The explosion of interest in CFC has led to the growing concern that existing methods for analysis may be prone to error and misinterpretation. In a recent article, Aru and colleagues [19] point out that existing cross-frequency coupling analyses are very sensitive to frequency band selection, noise, sharp signal transitions, and signal nonlinearities. Depending on the scenario, true underlying CFC can be missed, or spurious coupling can be detected. In addition, cross-frequency coupling methods tend to be statistically inefficient, requiring substantial amounts of data, making them unsuitable for time-varying scenarios or real-time application. Finally, in the absence of an appropriate statistical model, analysts typically employ surrogate data methods for statistical inference on cross-frequency coupling, making it difficult to properly answer even basic questions about the nature of the coupling, such as the size of the effect or its confidence/credible interval.

We describe here a novel method to estimate PAC that addresses these problems. A major source of error in existing methods stems from their reliance on traditional bandpass filtering. These filters can remove meaningful oscillatory coupling components (i.e., sidebands), and introduce spurious transients that resemble cross-frequency coupling. In our approach we use a state space oscillator model to separate out the different oscillations of interest. These models can preserve the relevant coupling terms in the signal and are resilient to noise and sharp signal transitions. We choose a particular model formulation, ingeniously proposed by Matsuda and Komaki, [20] that makes it straightforward to estimate both the phase and amplitude of oscillatory components. To further improve statistical efficiency, we introduce a parametric representation of the cross-frequency coupling relationship. A constrained linear regression estimates modulation parameters which can in addition be incorporated into a second state space model representing time-varying changes in the modulation parameters. Finally, we combine these statistical models to compute credible intervals for the observed coupling via resampling from the estimated posterior distributions. We demonstrate the efficacy of this method using simulated data, rat LFP data, and human EEG data.

We show that our method accurately estimates the parameters describing the oscillatory and modulation dynamics, provides improved temporal resolution, statistical efficiency, and inference compared to existing methods. Furthermore, we show that it overcomes the common problems with existing PAC methods described earlier, namely, band selection and spurious coupling introduced by sharp signal transitions and nonlinearities. The improved performance and robustness to artifacts should help improve the efficiency and reliability of PAC methods, and could enable novel experimental studies of PAC as well as novel medical applications.

## 2 Results

### 2.1 Overview of the State-Space PAC (SSP) method

In the conventional approach to phase and amplitude estimation, the signal is bandpass filtered to estimate the slow and fast components. The Hilbert transform is then applied to synthesize their imaginary counterparts. Finally, the slow component phase and the fast component amplitude are computed and used to calculate a Phase Amplitude Coupling (PAC) metric. In our approach, we use a state space model to estimate the oscillatory components of the signal, using the oscillation decomposition framework described by Matsuda and Komaki [20]. We assume, for the moment, that the observed signal *y*_*t*_ ∈ ℝ^2^ is a linear combination of two latent states representing a slow and a fast component 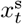 and 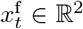, respectively. Each of these 2 dimensional latent states are assumed to be independesnt and their evolution over a fixed step size is modeled as a scaled and noisy rotation, for *j* = s, f:

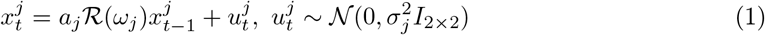

where *a*_*j*_ is a scaling parameter, 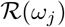 a 2-dimensional rotation of angle *ω*_*j*_ (the radial frequency) and 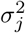 the process noise variance. An example of this state space oscillation decomposition is shown in Fig.1.a-d. This approach eliminates the need for traditional bandpass filtering since the slow and fast components are directly estimated under the model. Perhaps more importantly, the oscillations’ respective components can be regarded as the real and imaginary terms of a phasor or analytic signal. As a result, the Hilbert transform is no longer needed. Thus the latent vector’s polar coordinates provide a direct representation of the slow instantaneous phase 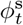 and fast oscillation amplitude 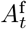 (Fig.1.f-g). We note 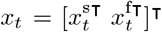 and obtain a canonical state space representation [21]:

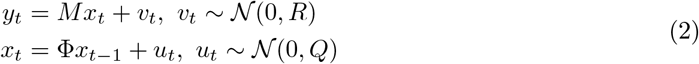

where Φ ∈ ℝ^4×4^ is a block diagonal matrix composed of the rotations described earlier, *Q* the total process noise covariance, *R* the observation noise covariance and *M* ∈ ℝ^1×4^ links the observation with the oscillation first coordinate. We estimate (Φ, *Q*, *R*) using an Expectation-Maximization (EM) procedure. To extend this model to add more independent oscillation components or to account for the harmonics of an oscillation (as one might encounter in a nonlinear system) we describe and derive a more general formulation in the Supplementary Materials 9.2.

**Figure 1:**
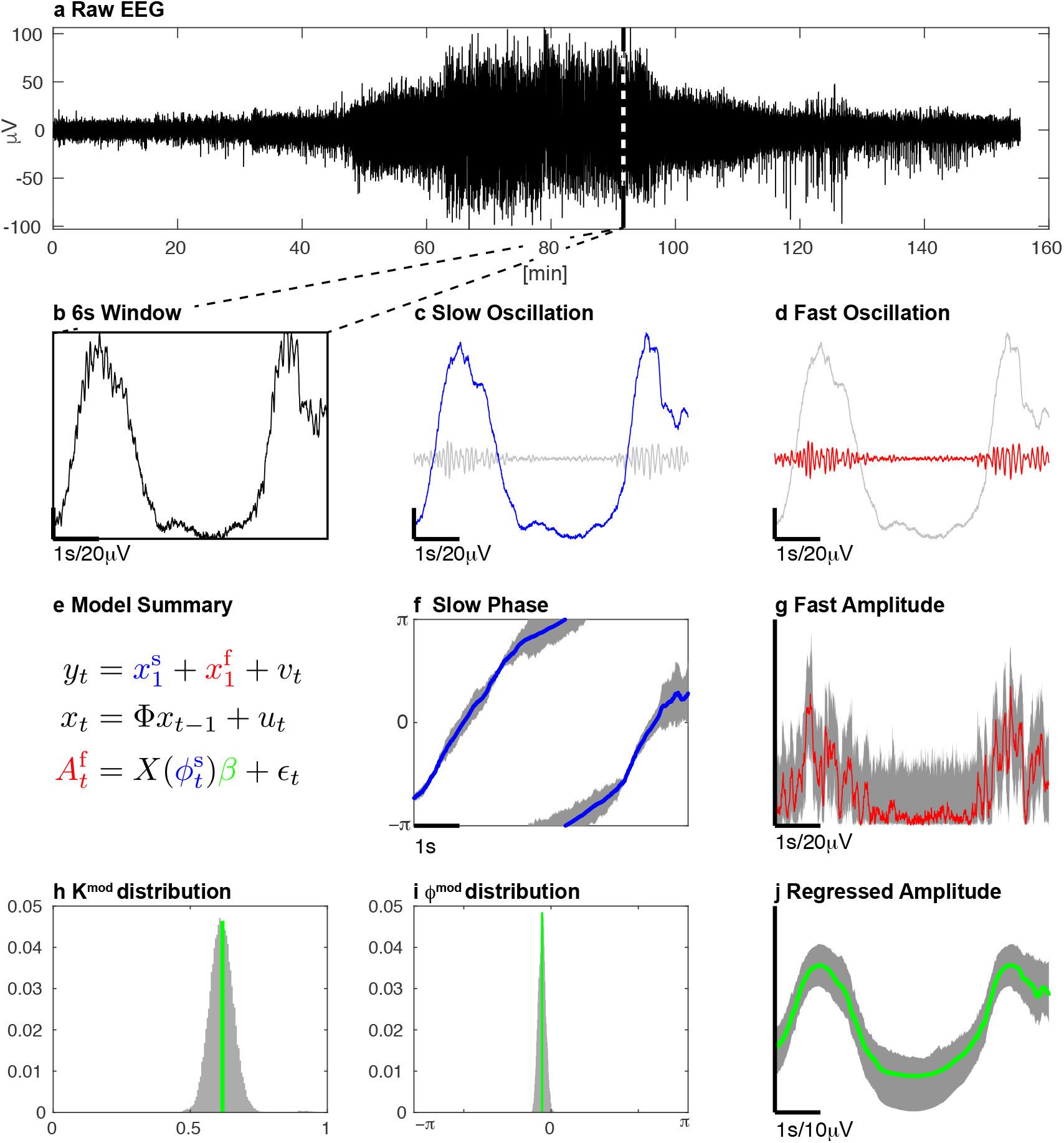
The oscillation decomposition for an EEG time-series from a human subject during anesthesia-induced unconsciousness using propofol. From the raw EEG trace (a), we extract a 6 second window (b) and decompose it into a slow (c) and a fast (d) oscillation using our state space model (e). We then deduce the slow oscillation phase (f) and the fast oscillation amplitude (g). Finally, we use a linear model (e) to regress the alpha amplitude (j) and to the estimate modulation parameters (h,j) and their distributions. Here, we used 200 × 200 resampled series (dark grey) to compute the 95% CI.

The standard approach for PAC analysis uses binned histograms to quantify the relationship between phase and amplitude [22] which is a major source of statistical inefficiency. Instead, we introduce a parametric representation of PAC based on a simple amplitude modulation model used in radio communications. To do so, we consider a constrained linear regression problem of the form 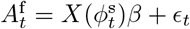 which can ultimately be rewritten:

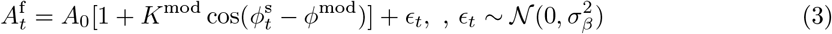

*K*^mod^ controls the strength of the modulation while *ϕ*^mod^ is the preferred phase around which the amplitude of the fast oscillation 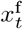 is maximal (Fig.1.h-j). For example, if *K*^mod^ = 1 and *ϕ*^mod^ = 0, the fast oscillation is strongest at the peak of the slow oscillation. On the other hand, if *ϕ*^mod^ = *π*, the fast oscillation is strongest at the trough or nadir of the slow oscillation.

Finally, instead of relying on surrogate data [19] to determine statistical significance, which decreases efficiency even further, our model formulation allows us to estimate the posterior distribution of the modulation parameters *p*(*K*^mod^, *ϕ*^mod^|{*y*_*t*_}_*t*_) and to deduce the associated credible intervals (CI) (Fig.1.f-j).

We refer to our approach as the State-Space PAC (SSP) method. Because physiological systems are time varying, we apply it over multiple non-overlapping windows. In a variation of our method, we can also impose a temporal continuity constraint on the modulation parameters across windows using a second state-space model, yielding what we term the double State Space PAC estimate (dSSP).

### 2.2 Human EEG Data

To demonstrate the performance of our methods we first analyzed EEG data from a human volunteer receiving propofol to induce sedation and unconsciousness (Fig. 2). As expected, as the concentration of propofol increases, the subject’s probability of response to auditory stimuli decreases. The parametric power spectral density (see equation (38), in Suplementary material 9.1) changes during this time, developing beta (12.5-25 Hz) oscillations as the probability of response begins to decrease, followed by slow (0.1-1 Hz) and alpha (8-12 Hz) oscillations when the probability of response is zero (Fig. 2-d) as in [23]. For a window *T*, we estimate the modulation strength 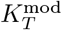 and phase 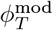 (and CI) with dSSP (Fig. 2-f) and we gather those estimates in the Phase Amplitude Modulogam: PAM(*T*, *ψ*) (Fig. 2-e). For a given window *T*, PAM(*T*,.) is a probability density function (pdf) having support [−*π*, *π*]. It assesses how the amplitude of the fast oscillation is distributed with respect to the phase of the slow oscillation. When the probability of response is zero, we observe a strong “peak-max” 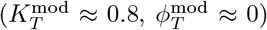 pattern in which the fast oscillation amplitude is largest at the peaks of the slow oscillation. During the transitions to and from unresponsiveness, we observe a “trough-max” pattern of weaker strength (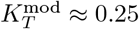, 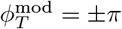) in which the fast oscillation amplitude is largest at the troughs of the slow oscillation. Note that the coefficient of determination *R*^2^ for the modulation relationship mirrors the coupling strength *K*^mod^ since 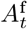 is better predicted by our model when the coupling is strong.

**Figure 2:**
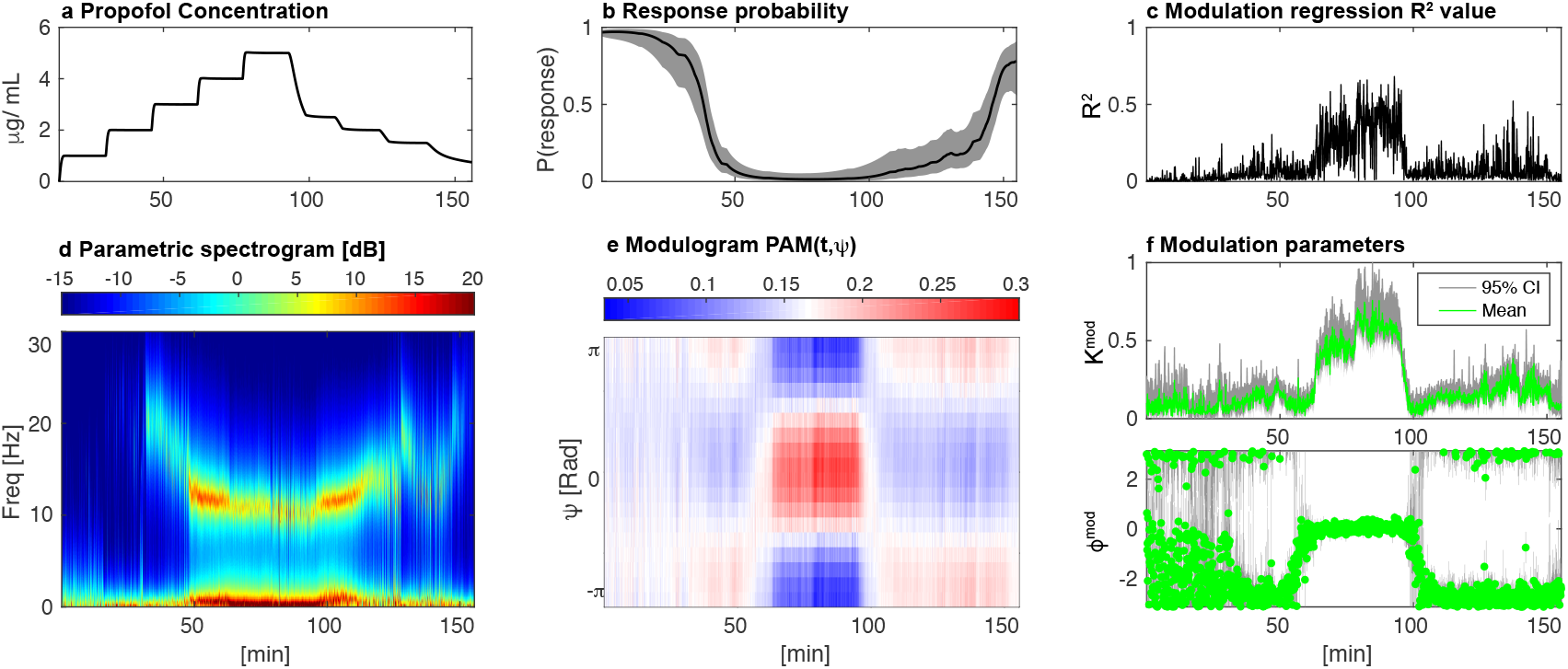
Propofol-induced unconsciousness in a human subject monitored with EEG. Increasing target effect-site concentrations of propofol were infused (a) while loss and recovery of consciousness were monitored behaviorally with an auditory task from which a probability of response was estimated (b). *R*^2^ value of our modulation regression (c). dSSP was used to estimate the parametric spectrogram (d), the phase amplitude modulogram (e) and the modulation parameters (f) *K*^mod^ and *ϕ*^mod^ alongside their CI computed with 200 × 200 samples.

When averaged over long, continuous and stationary time windows, conventional methods provide good qualitative assessments of PAC. However, in many cases, analyses over shorter windows of time may be necessary if the experimental conditions or clinical situation changes rapidly. In previous work [23], we analyzed PAC using conventional methods with relatively long *δt* = 120*s* windows, appropriate in this case because propofol was administered at fixed rates over ~ 14 minute intervals. The increased statistical efficiency of the SSP and dSSP methods makes it possible to analyze much shorter time windows of *δt* = 6*s*, which we illustrate in two subjects, one with strong coupling (Fig. 3) and another with weak coupling (Supplementary Fig. 10). To do so, we compare SSP, dSSP and standard methods used with *δt* = 120*s* or *δt* = 6*s* based on the modulogram and on the Modulation Index (MI) estimates. The latter assesses the strength of the modulation by measuring, for any window *T* how different PAM(*T*,.) is from the uniform distribution. The Kullback-Leibler Divergence is typically used for this purpose. Thus, any random fluctuations in the estimated PAM will increase MI, introducing a bias. Our model parametrization is used to derive PAM, MI and associated CI but standard non-parametric analysis typically rely on binned histogram. As a results they estimate statistical significance by constructing surrogate datasets and reporting p-values [19].

**Figure 3:**
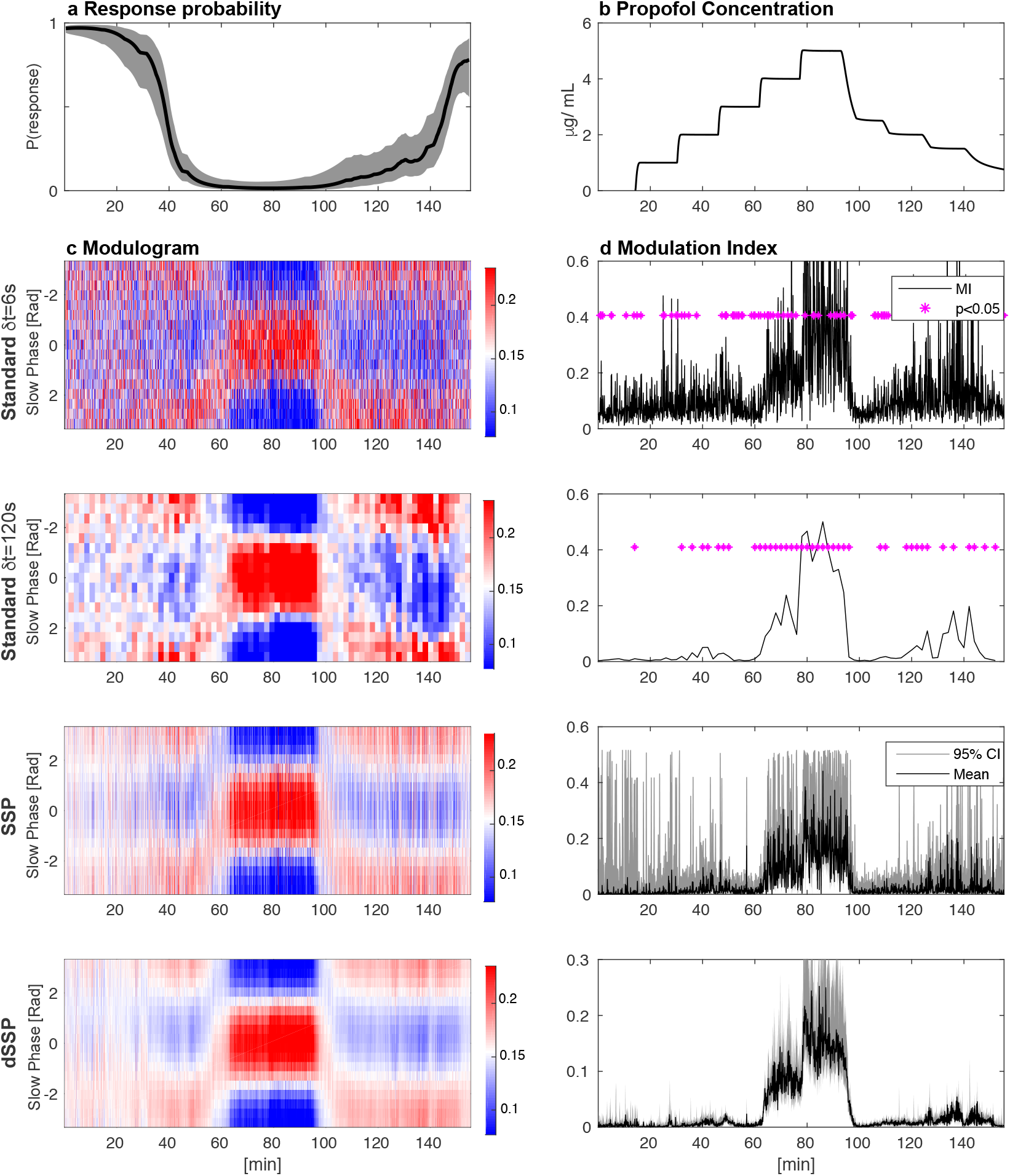
The Phase Amplitude Coupling profile of a subject infused with increasing target effect site concentrations of propofol. Left: response probability curves (a) aligned with modulograms (c) (distribution of alpha amplitude with respect to slow phase) computed with standard (top) and state-space parametric (bottom) methods. Right: propofol infusion target concentration (b) aligned with corresponding Modulation Indices (d). Standard technique significance was assessed using 200 random permutations and CI where estimated using 200 × 200 samples

Both subjects exhibit the typical phase amplitude modulation profile previously described when they transition in and out of unconsciousness. Nevertheless, since SSP more efficiently estimates phase and amplitude [20] and produces smooth PAM estimates even on short windows, MI estimates derived from SSP show less bias that the standard approach. For the same reasons, *ϕ*^mod^ estimates show less variance than the standard approach. The dSSP algorithm provides a temporal continuity constraint on the PAM, making it possible to track time-varying changes in PAC while further reducing the variance of the PAM estimates. Finally, our parametric strategy provides posterior distributions for *K*^mod^, *ϕ*^mod^ and MI, making it possible to estimate CI for each variable and it assesses significance without resorting to surrogate data methods.

### 2.3 Rat LFP Data

To illustrate the performance of our approach in a different scenario representative of invasive recordings in animal models, we analyzed rat LFP during a learning task hypothesized to involve theta (6-10Hz) and low gamma (25-60Hz) oscillations. We applied dSSP on 2 second windows (Fig. 4) and confirmed that theta-gamma coupling in the CA3 region of the hippocampus increases as the rat learned the discrimination task, as originally reported in Tort et al. [4]. In our analysis using dSSP, we did not pre-select the frequencies of interest, nor did we specify bandpass filtering cutoff frequencies. Rather, the EM algorithm was able to estimate the specific underlying oscillatory frequencies for phase and amplitude from the data, given an initial starting point in the theta and gamma ranges. Thus we illustrate that our method can be applied effectively to analyze LFP data, and that it can identify the underlying oscillatory structure without having to specify fixed frequencies or frequency ranges.

**Figure 4:**
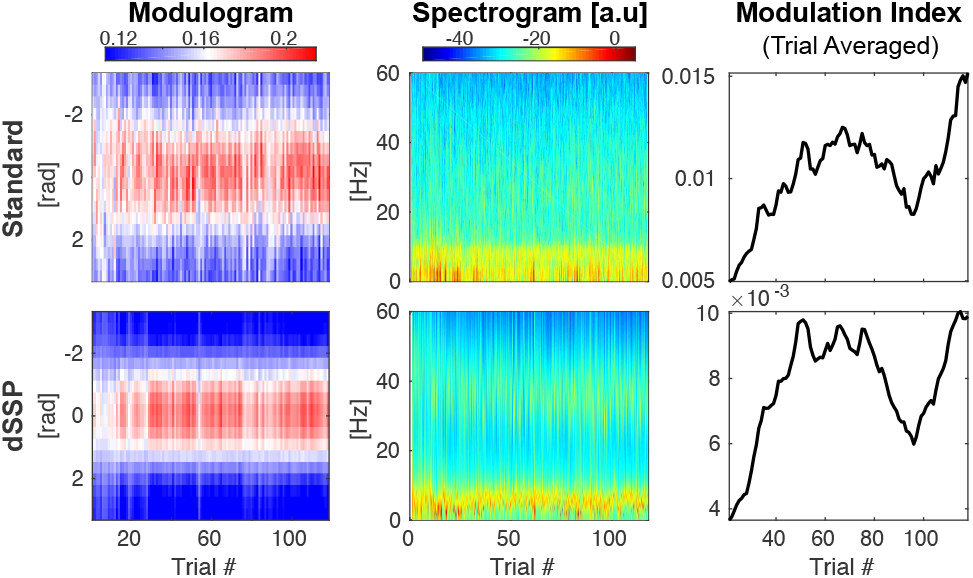
Rats show increased theta-gamma coupling when learning a discrimination task in hypocampus CA3 region. Top: Standard [4] processing. Bottom: dSSP applied on 2 second windows.

### 2.4 Simulation Studies

To test our algorithms in a more systematic way as a function of different modulation features and signal to noise levels, we analyzed multiple simulated data sets. By design, these simulated data were constructed using generative processes or models different than the state space oscillator model; i.e., the simulated data generating processes were outside the “model class” used in our methods. Here, we focus on slow and an alpha components to reproduce our main experimental data cases. In doing so, our intent is not to provide an exhaustive characterization of the precision and accuracy of our algorithm, since this would strongly depend on the signal to noise ratio, the signal shape, etc. Instead, we aim to illustrate how and why our algorithm outperforms standard analyses in the case of short and noisy time-varying data sets.

We first compare the resolution and robustness of dSSP with conventional techniques on broad-band signals with modulation parameters varying on multiple time scales. Results are reported for different generative parameters (See Section 8.3.2, 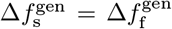,*σ*_s_ and *σ*_f_) in Fig. 5 and Supplementary Fig. 11 and associated signal traces are illustrated Supplementary Fig. 12. Although robust when averaged on long windows with stationary coupling parameters, standard techniques become ineffective when the modulation parameters vary rapidly across windows. The modulation cannot be resolved when long windows are used. However if we reduce the window size to compensate, the variance of the estimates increases significantly. A trade-off has to be found empirically. On the other hand, we see that, applied on 6-second windows, (d)SSP can track the rapid changes in amplitude modulation even in the case of a low signal to noise ratio. The dSSP algorithm also provides estimates of the posterior distribution of the modulation parameters, making it straightforward to construct CI and perform statistical inference. By comparison, the surrogate data approach becomes infeasible as there are fewer and fewer data segments to shuffle.

**Figure 5:**
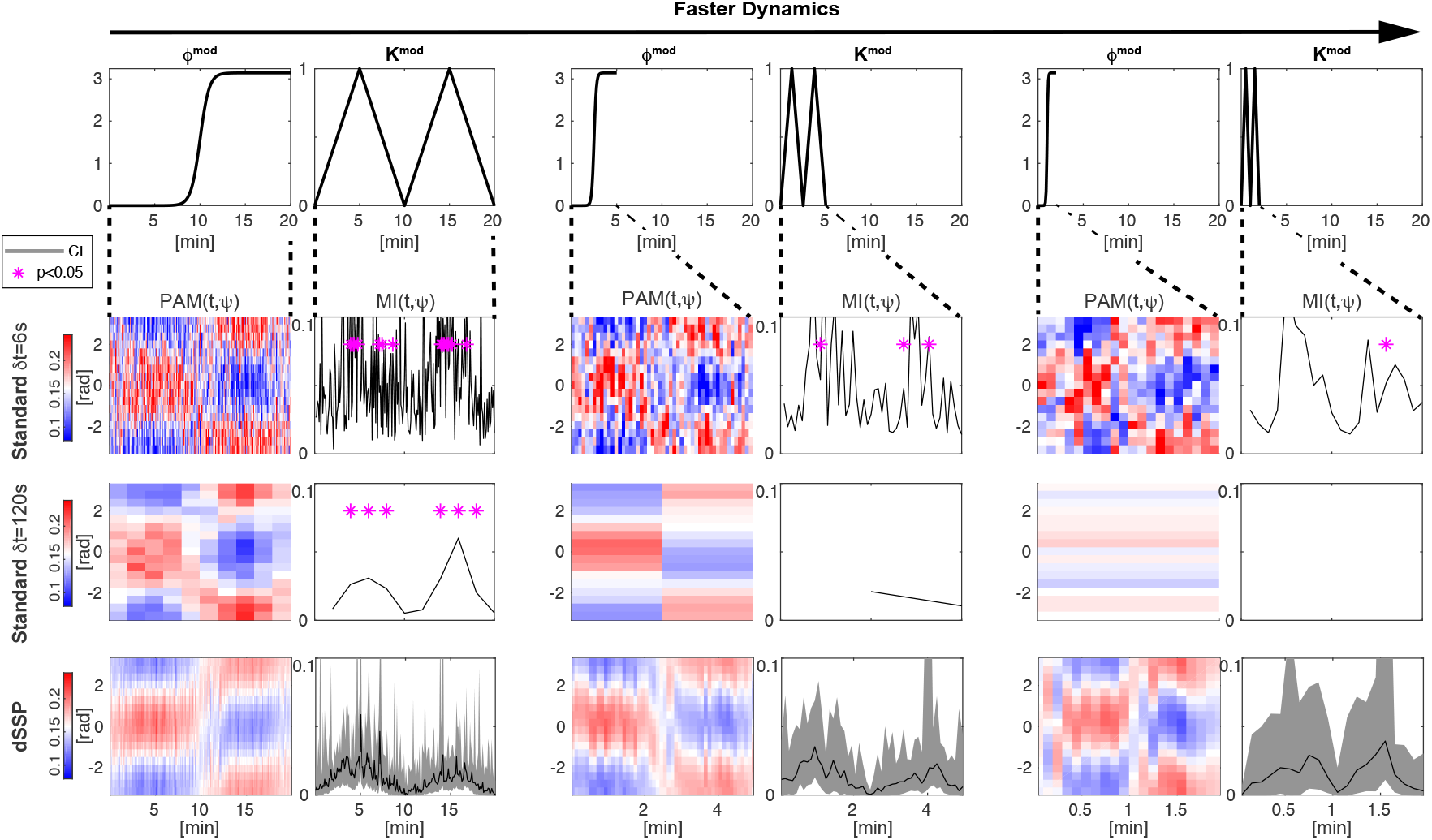
Comparison of the modulation estimates using standard methods and our new dSSP method. Slow and fast oscillations were generated by filtering white noise around *f*_s_ =1Hz and *f*_s_10Hz with 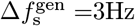 and normalized to standard deviation *σ*_s_ =0.5 and *σ*_f_ =2. The time scale over which *K*^mod^ and *ϕ*^mod^ changed varied between 20 minutes to 5 and 2 minutes. See Fig. 12 for typical signal traces.

In a recent paper, Dupré la Tour et al. [24] designed an elegant nonlinear PAC formulation, described as a driven autoregressive (DAR) process, where the modulated signal is a polynomial function of the slow oscillation. The latter, referred to as the driver, is filtered out from the observation around a preset frequency and used to estimate DAR coefficients. The signal parametric spectral density is subsequently derived as a function of the slow oscillation. The modulation is then represented in terms of the phase around which the fast oscillation power is preferentially distributed. A gridsearch is performed on the driver, yielding modulograms for each slow central frequency over a range of fast frequencies. The frequencies associated with the highest likelihood and/or strongest coupling relationship are then selected as the final coupling estimate.

This parametric representation improves efficiency, especially in the case of short signal windows, but because it relies on an initial filtering step, it also shares some of the limitations of conventional techniques. As we will see, spurious CFC can emerge from abruptly varying signals or nonlinearities. Additionally, this initial filtering step might contaminate PAC estimates from short data segments with wideband slow oscillations.

To compare our methods with standard techniques and the DAR method, we generated modulated signals with the scheme described in Dupré la Tour et al. [24] (equation (26), *λ* = 3, and *ϕ*^mod^ = −*π*/3) using different frequencies of interest (*f*_s_ and *f*_f_) spectral widths 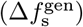 and Signal to Noise Ratios (SNR). Typical signal traces for those generating parameters are reported in Supplementary Fig. 14 and 15. We then compare how well these methods recover the slow oscillation and the fast oscillation (Supplementary Fig. 14 and 15) or the modulation phase (Fig. 6 and Supplementary Fig.13). Contrary to the other methods presented here, SSP does not compute the full comodulograms to select frequencies of interest but rather identifies them by fitting the state space oscillator model. Coupling is only estimated in a second step. Although we used tangible prior knowledge in previous sections to initialize the algorithm, we adapt an initialization procedure from [25] (See Supplementary Materials 9.6) to provide a fair comparison. For each condition, we generated 400 six-second windows. When necessary, the driver was extracted using 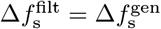.

**Figure 6:**
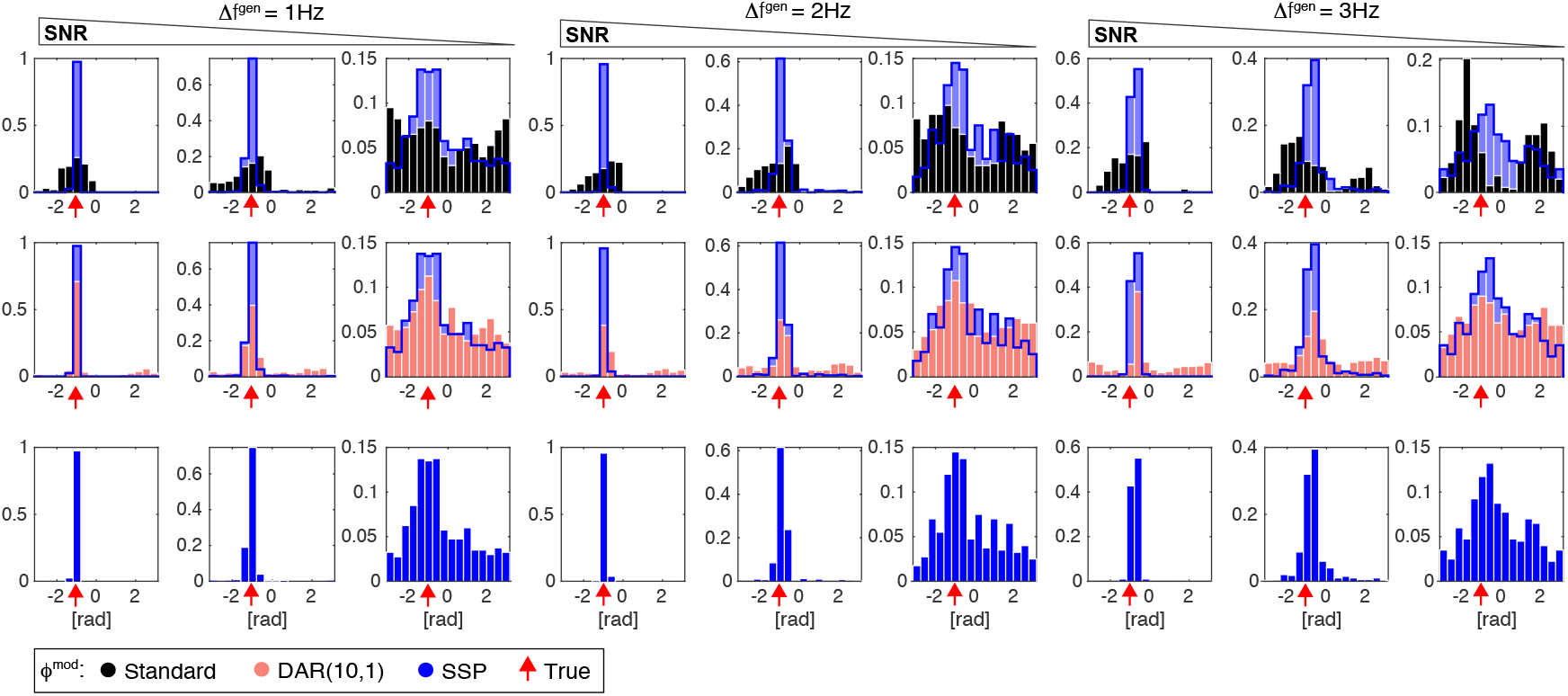
Modulation phase *ϕ*^mod^ estimation and comparisons with standard methods (black), DAR (pink) and SSP (blue). 400 windows of 6 seconds were generated with: a slow oscillation (filtered from white noise around *f*_s_ = 1Hz with bandwidth 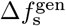, normalized to standard deviation *σ*_s_) and a modulated fast oscillation (*ϕ*^mod^ = −*π*/3, modeled with a sinusoid *f*_s_ = 10Hz and normalized to *σ*_f_). We added unit normalized Gaussian noise and we used 3 Signal To Noise Ratio (SNR) conditions ((*σ*_s_, *σ*_s_) = (2, 1.5), (1, 0.6) and (0.7, 0.3)). We show typical signal traces for these different conditions in Supplementary Materials Fig 14.

We find that our algorithm better retrieves fast frequencies in each case (Supplementary Fig. 14 and 15) especially when the slow oscillation is wider-band. It also outperforms the other methods when estimating modulation phase (Fig. 6 and Supplementary Fig.13): our algorithm is stable in the case of broadband 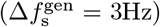 or weak ((*σ*_s_, *σ*_s_) = (0.7, 0.3)) slow oscillations and *ϕ*^mod^ is estimated accurately with very few outliers and a smaller standard deviation in virtually all cases considered.

### 2.5 Overcoming Key Limitations of CFC Analysis: Sharp Transitions, Nonlinearities, and Frequency Band Selection

Despite the central role that CFC likely plays in coordinating neural systems, standard methods of CFC analysis are subject to many caveats that are a source of ongoing concern [19]. Spurious coupling can arise when the underlying signals have sharp transitions or nonlinearities. On the other hand, true underlying coupling can be missed if the frequency band for bandpass filtering is not selected properly. Here we illustrate how our SSP method is robust to all of these limitations. We also show how our method is able, counterintuitively, to extract nonlinear features of a signal using a linear model.

#### 2.5.1 Signals with abrupt changes and/or harmonics

Oscillatory neural waveforms may have features such as abrupt changes or asymmetries that are not confined to narrow bands [26]. In such cases, truncating their spectral content with standard bandpass filters can distort the shape of the signal and can introduce artefactual components that may be incorrectly interpreted as coupling.

The state space oscillator model provides an alternative to bandpass filtering that can accommodate non sinusoidal wave-shapes. In this section, we extend the model to explicitly represent the slow oscillatory signal’s harmonics, thus allowing the model to better represent oscillations that have sharp transitions and those that may be generated by nonlinear systems. To do so, we optimize *h* oscillations with respect to the same fundamental frequency *f*_s_ (see Supplementary Materials 9.2.3). We further combine this model with information criteria (Akaike Information Criteria -AIC- [27] or Bayesian Information Criteria -BIC- [28]) to determine (i) the number of slow harmonics *h* and (ii) the presence or the absence of a fast oscillation. We select the best model by minimizing ΔIC = IC − min(IC). We only report AIC based PAC estimation here although both AIC and BIC perform similarly. When multiple slow harmonics are favored, we use the phase of the fundamental oscillation to estimate PAC.

We first simulated a non symmetric abruptly varying signal using a Van der Pol oscillator (equation (27), *ϵ* = 5, *ω* = 5*s*^−1^) to which we added observation noise 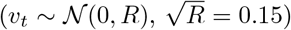. We then considered two scenarios, one with a modulated fast sinusoidal wave (Fig. 7.a, 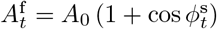,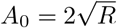 and *f*_f_ = 10Hz), and one without (Fig. 7.b). Because our model is able to fit the sharp transitions, both AIC and BIC identify the correct number of independent components (Fig. 7.a-4 and b-4: orange and grey traces relative position is switched when adding or not a fast oscillation). As a consequence, when no clear fast oscillation is detected, no PAC is calculated (Fig. 7.a-6). On the other hand, when no fast oscillation is present, standard techniques extract a fast component stemming from the abruptly changing slow oscillation, leading to the detection of spurious coupling (Fig. 7.a-3).

**Figure 7:**
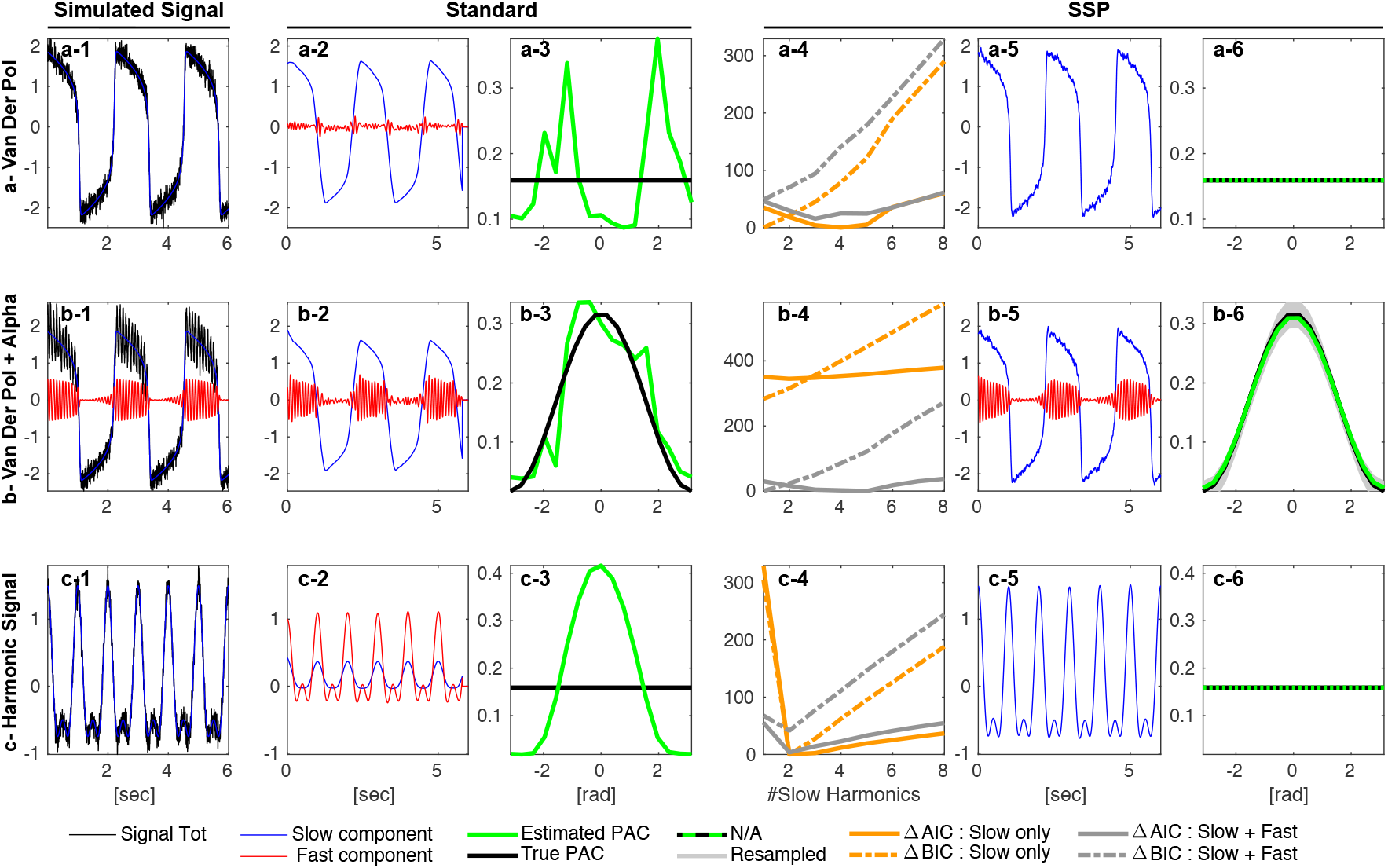
PAC analysis of 6-second signals with harmonic content using standard methods and SSP. The signal was either generated using a Van der Pol oscillator alone (a-1), a Van der Pol oscillator with a modulated alpha oscillation (b-1), or with a nonlinearity according to equation (4)(c-1). The standard method use conventional filters to extract the oscillation (0.1-1Hz and 6-14Hz (ab-2) and 0.6-1.2Hz and 0.9-3.1Hz (c-2)). SSP was combined with an information criteria (AIC or BIC)(abc-4) to select the optimal number of independent oscillations (one or two) and the number of slow harmonics (abc-5). PAC is reported as the distribution of the fast amplitude with respect to the slow phase (abc-3 and abc-6). For SSP 200 samples were drawn from the posterior to generate CI (b-6).

Nonlinear inputs arising from signal transduction harmonics are a similar hurdle in CFC analysis. If we consider a slow oscillation 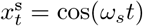 non-linearly transduced as 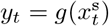, we can write a second order approximation:

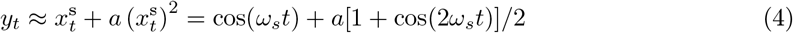

If *ω*_s_/(2*π*) = 1Hz, bandpass filtering *y*_*t*_ a round 0.9 − 3.1Hz to extract an oscillation peaking at *f*_f_ = 2Hz would yield [19]:

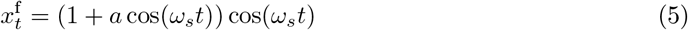

In such a case, standard CFC analysis infers significant coupling (Fig. 7.c-3) while oscillation decomposition correctly identifies a harmonic decomposition without CFC (Fig. 7.c-6).

This model selection strategy does not guarantee that the correct model will always be selected. Furthermore, the oscillation decomposition itself is often a non convex optimization problem. However, we observe that the (extended) state-space oscillator is better suited to model physiological signals than narrow band components. In addition, the model selection paradigm combined with prior knowledge of the signal content (e.g., propofol anesthesia slow-alpha or rodent hippocampal theta-gamma oscillations) allows us to study PAC in a more principled way.

#### 2.5.2 Frequency Band Selection

If bandpass filters with an excessively narrow bandwidth are applied to a modulated signal, the modulation structure can be obliterated. Let us consider the following signal:

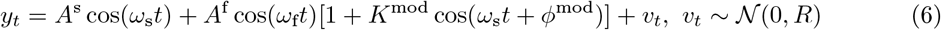

Developing *y*_*t*_ yields 4 frequency peaks: the slow and fast frequencies *ω*_s_ and *ω*_f_ and two sidelobes centered around *ω*_f_ − *w*_s_ and *ω*_f_ + *ω*_s_:

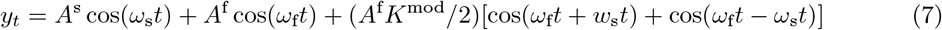

As a consequence, if the fast oscillation is extracted without its side lobes, no modulation is detected, as illustrated Fig. 8-c.

**Figure 8:**
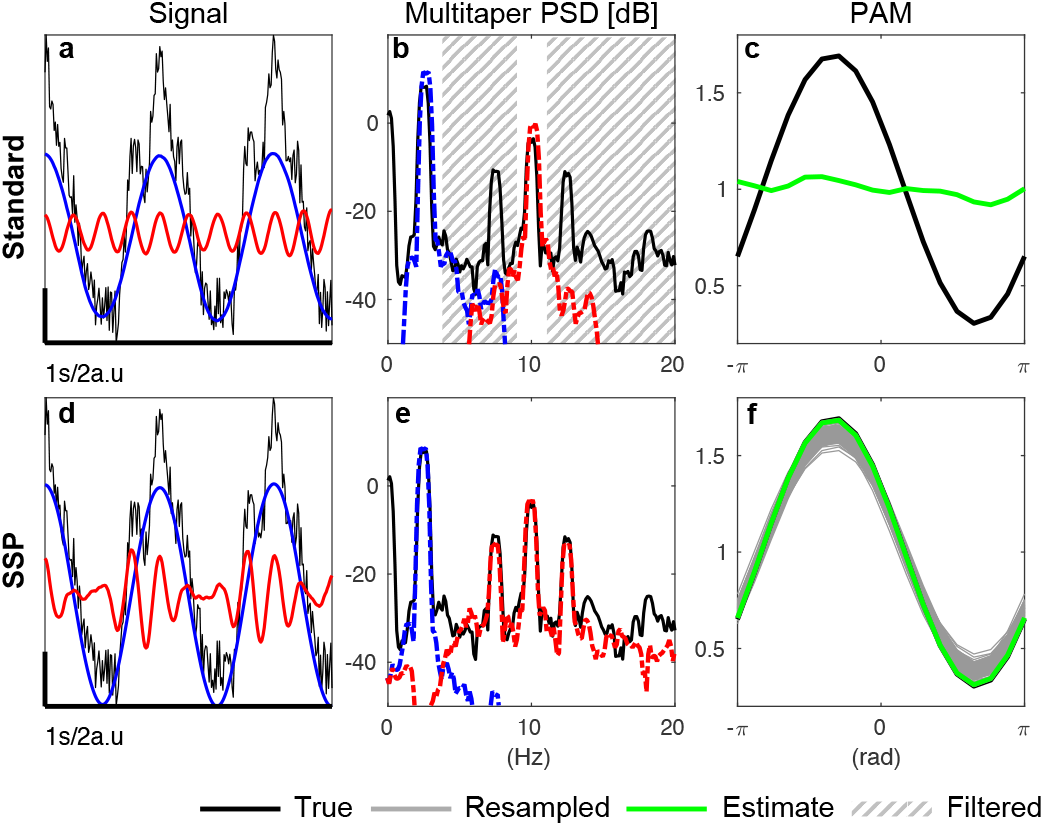
Decomposition (a,d), power spectral density (b,e) and modulogram (c,f). The top row shows the result of applying a narrow bandpass filter that removes the modulation s ide lobes. The bottom row shows the result of applying the oscillation decomposition used in SSP and dSSP, which preserves the modulation structure. (*K*^mod^ = 0.6, *ϕ*^mod^ = −π/3, *R* = 4, 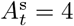 and 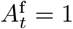)

Our SSP algorithm uses a state-space oscillator decomposition which does not explicitly model the structural relationship giving rise to the modulation side lobes (equation (6)). Yet, we see that the modulation is successfully extracted, as observed in the fitted time series (Fig. 1) and in the spectra (Fig. 8-e). The model is able to achieve this by making the frequency response of the fast component wide enough to encompass the side lobes. The algorithm does this by inflating the noises covariances *R* and 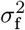 and 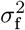 and deflating *a*_f_. In theory it might be possible to use a higher order model like an ARMA(4,2) (which would represent the product of 2 oscillations and which poles are in 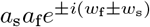, or to directly model coupling through a nonlinear observation. However, in both cases, we found that such models were difficult to fit to the data, and quickly became underconstrained when applied to noisy, non-stationary, non-sinusoidal physiological signals. Instead, we found that our simpler model was able to pull out the modulated high-frequency component robustly.

In summary, the first stage of our algorithm can extract nonlinearities stemming from the modulation before fitting them with a regression model in the second stage. The main consequence of this approach is to inflate the variance in the fast component estimation. See for example the wide CI in the fast oscillation estimate in Fig. 1g. In turn, we resample the fast oscillation amplitudes from a wider distribution than is actually the case. Although this does not affect the estimates of *ϕ*^mod^, it does produce a conservative estimate when resampling *K*^mod^, i.e., the credible intervals are wider than they might be otherwise. Even so, we find that our approach still performs better than previous methods (Fig. 5,3 and 6).

## 3 Discussion

We have presented a novel method that integrates a state space model of oscillations with a parametric formulation of phase amplitude coupling (PAC). Under this state space model we represent each oscillation as an analytic signal [20] to directly estimate the phase or amplitude. We then characterize the PAC relationship using a parametric model with a constrained linear regression. The regression coefficients, which efficiently represent the coupling relationship in only a few parameters, can be incorporated into a second state space model to track time-varying changes in the PAC. We demonstrated the efficacy of this method by analyzing neural time series data from a number of applications, and illustrated its improved statistical efficiency compared to standard techniques using simulation studies based on different generative models. Finally, we showed how our method is robust to many of the limitations associated with standard phase amplitude coupling analysis methods.

The efficacy of our method stems from a number of factors. First, the state-space analytic signal model provides direct access to the phase and amplitude of the oscillations being analyzed. This linear model also has the remarkable ability to extract a nonlinear feature (the modulation) by imposing “soft” frequency band limits which are estimated from the data. The oscillation decomposition is thus well-suited to analyze physiological signals that are not confined to strict band limits. We also proposed a harmonic extension that can represent nonlinear oscillations (e.g., Van der Pol, Fig. 7), making it possible to better differentiate between true and spurious PAC resulting from bandpass filtering artifacts. The parametric representation of the coupling relationship can accommodate different modulation shapes and increases the model efficiency even further.

Overall, we addressed a majority of the significant limitations associated with current methods for PAC analysis. The neural time series are processed more efficiently (Fig. 3), frequency bands of interest are automatically selected (Fig. 4), extracted (Fig. 8.d-e), and more realistically modeled (Fig. 7). Contrary to standard methods, we do not need to average PAC-related quantities across time, reducing the amount of contiguous time series data required. Moreover, the posterior distributions of the signals of interest are well-defined under our proposed model. Sampling from them bypasses the need to build surrogate data, which can obscure non-stationary structure in the data and underestimate the false positive rate [19]. Conversely, because SSP estimates the modulation parameters’ posterior distribution, we report CI and provide information on the statistical significance of our results as well as the strength and direction of the modulation. Our dynamic estimation of PAC hence makes it possible to base inference on much shorter windows -as short as 6 seconds for slow 0.1-1Hz signals. Other novel models have been proposed to represent PAC, including driven autoregressive models (DAR) [24] and generalized linear models (GLM) [29]. As we saw earlier, SSP performs better than the DAR and standard approaches, particularly when the signal to noise is low. The GLM represents the modulation relationship parametrically as we do, but in a more general form, and provides confidence intervals using the bootstrap [29]. Both the DAR and GLM approaches remain reliant on traditional bandpass filtering for signal extraction, and thus remain vulnerable to the crucial problems introduced by these filters [19]. Our method is the first to use state space models combined with a parametric model of the modulation, the latter of which could be generalized in the manner described by [29]

Such improvements could significantly improve the analysis of future studies involving CFC, and could enable medical applications requiring near real-time tracking of CFC. One such application could be EEG-based monitoring of anesthesia-induced unconsciousness. During propofol induced anesthesia, frequency bands are not only very well defined, but the PAC signatures strongly discriminates deep unresponsiveness (peak-max) from transition states (through-max), most likely reflecting underlying changes in the polarization level of the thalamus [30]. Thus, PAC could provide a sensitive and physiologically-plausible marker of anesthesia-induced brain states, offering more information than spectral features alone. Accordingly, a recent study [31] reported cases in which spectral features could not perfectly predict unconsciousness in patients receiving general anesthesia. In this same data, CFC measures (peak-max) could more accurately indicate a fully unconscious state from which patients cannot not be aroused [32]. In the operating room, drugs can be administered rapidly through bolus injection, drug infusion rates can change abruptly, and patients may be aroused by surgical stimuli, leading to corresponding changes in patients’ brain states over a time scale of seconds [33] [34]. These rapid transitions in state can blur modulation patterns estimated using conventional methods. Faster and more reliable modulation analysis could therefore lead to tremendous improvement in managing general anesthesia. Conventional methods are impractical since they require minutes of data to produce one estimate; in contrast our method can estimate CFC on a time-scale compatible with such applications.

Since CFC analysis methods were first introduced into neuroscience, there has been a wealth of data suggesting that CFC is a fundamental mechanism for brain coordination in both health and disease [15] [35]. Our method addresses many of the challenging problems encountered with existing techniques, while also significantly improving statistical efficiency and temporal resolution. This improved performance could pave the way for important new discoveries that have been limited by inefficient analysis methods, and could enhance the reliability and efficiency of PAC analysis to enable their use in medical applications.

## 4 Acknowledgements

This work was generously supported by the Bertarelli Foundation (fellowship for H.S.), the National Science Foundation (predoctoral fellowship for A.M.B.), and grants from the National Institutes of Health (P01GM118269, R01AG056015, R01AG054081, R21DA048323 to P.L.P.). Acknowledgements. We thank Adriano Tort and colleagues for sharing their rat LFP dataset.

## 5 Author Contributions

Conceptualization: H.S., P.L.P. Methodology: H.S., E.P.S, A.M.B., P.L.P. Software H.S., E.P.S, A.M.B. Formal Analysis: H.S., P.L.P. Writing: H.S., P.L.P. Funding Acquisition: P.L.P.

## 7 Data materials and availability

Data and code will be made available in a public database prior to publication.

## 8 Methods

### 8.1 State-Space Oscillator Model

For a time series sampled at *F*_*s*_, we consider a time window 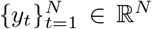, and we assume, in this section, that *y*_*t*_ is the sum of observation noise and components from two latent states 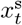 and 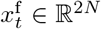 which account for a slow and a fast component.^1^ We use the oscillation decomposition model described by Matsuda and Komaki [20]. For *j* = s, f and *t* = 2‥*N*, each component follows the process equation:

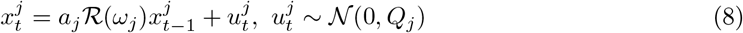

where *a*_*j*_ ∈ (0, 1) and 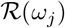 is a rotation matrix with angle *ω*_*j*_ = 2*πf*_*j*_/*F*_*s*_

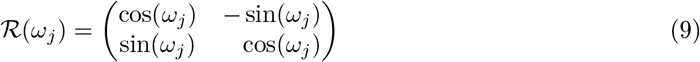

and

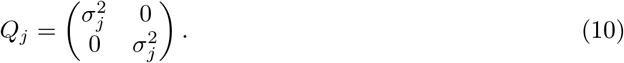

As previously stated, the phase 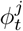 and amplitude 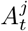 of each oscillation are obtained using the latent vector polar coordinates:

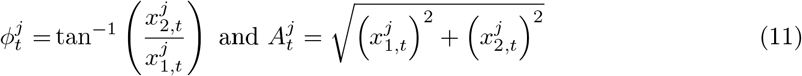

Each oscillation has a broad-band power spectral density (PSD) with a peak at frequency *f*_*j*_. The parametric expression for this PSD is derived in the Supplementary Materials 9.1.

Setting *M* = [1 0 1 0], 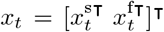, and *Q* and Φ to be block diagonal matrices whose blocks are *Q*_*j*_ and 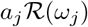, respectively, we find the canonical state space of equation (2).

Given the observed signal *y*_*t*_, we aim to estimate both the hidden oscillations *x*_*t*_ and their generating parameters (Φ, *Q*, *R*). We do so using an Expectation-Maximization (EM) algorithm (see Supplementary Materials 9.2 for a more general derivation). The hidden oscillations *x*_*t*_ are estimated in the E-step of the EM algorithm using the Kalman filter and fixed-interval smoother [49], while the generating parameters are estimated in each iteration of the M-step.

### 8.2 Phase Amplitude Coupling Model

#### 8.2.1 Standard Processing Using Bandpass Filters and the Hilbert Transform

Standard approaches for PAC analysis follow a procedure described in Tort, et al. [22], we briefly summarize here. The raw signal *y*_*t*_ is first bandpass filtered to isolate slow and fast oscillations. A Hilbert transform is then applied to_s_ estimate the instantaneous phase of the slow oscillation 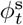, and instantaneous amplitude of the fast oscillation 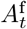. At time *t*, the alpha amplitude 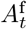 is assigned to one of (usually 18) equally spaced phase bins of length *δψ* based on the instantaneous value of the slow oscillation phase: 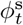. The histogram is constructed over some time window *T* of observations, for instance a ~ 2 minute epoch, which yields the phase amplitude modulogram (PAM) [38]:

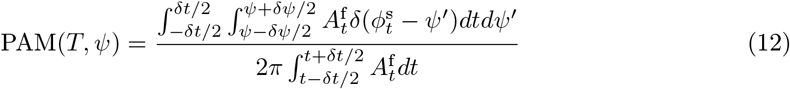

For a given window *T*, PAM(*T*,.) is a probability distribution function which assesses how the fast oscillation amplitude is distributed with respect to the slow oscillation phase. The strength of the modulation is then usually measured with the Kullback–Leibler divergence with a uniform distribution. It yields the Modulation Index (MI):

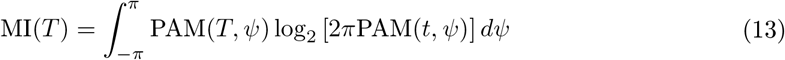

Finally, under this standard approach, surrogate data such as random permutations are used to assess the statistical significance of the observed MI. Random time shifts Δ*t* are drawn from a uniform distribution whose interval depends on the problem dynamics [38] and phase amplitude coupling is estimated using the shifted fast amplitudes 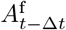 and the original slow phase 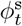. The MI is then calculated for this permuted time series, and the process is repeated to construct a null distribution for the MI. The original MI is deemed significant if it is bigger that 95% of the permuted values. Overall, this method requires that the underlying process remains stationary for sufficiently long so that the modulogram can be estimated reasonably well and so that enough comparable data segments can be permuted in order to assess significance.

#### 8.2.2 Parametric Phase Amplitude Coupling

To improve statistical efficiency, we introduce a parametric representation of PAC. For a given window, we consider the following (constrained) linear regression problem:

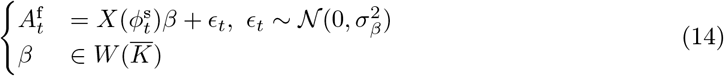

where *β* = [*β*_0_ *β*_1_ *β*_2_]^**T**^, 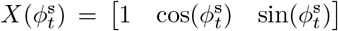 and 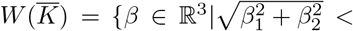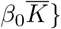. If we define:

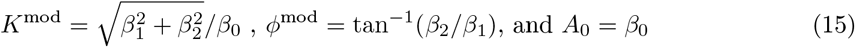

we see that equation (14) is equivalent to:

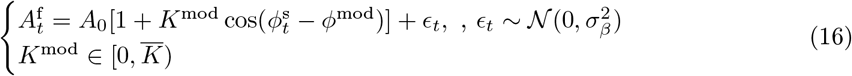

Setting 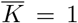 ensures that the model is consistent, i.e., that the modulation envelope cannot exceed the amplitude of the carrier signal. But this can be a computationally expensive constraint to impose. If the data have a high signal to noise ratio so that *K*^mod^ is unlikely to be greater than 1 by chance, we could also choose to solve the unconstrained problem 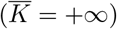. Under the constrained solution, the posterior distribution for *β* is a truncated multivariate t-distribution [45]:

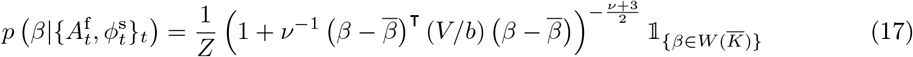

The likelihood, conjugate prior, posterior parameters 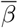, *V*, *b*, *v*, and the normalizing constant *Z* are justified and derived in Supplementary Material 9.3. We refer to this estimate as State Space PAC (SSP) and we note:

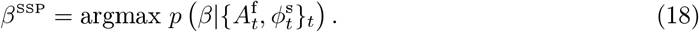

#### 8.2.3 Posterior sampling

The standard approach relies on surrogate data to determine statistical significance, which decreases its efficiency even further. Instead, we estimate the posterior distribution *p*(*β*|{*y*_*t*_}_*t*_) from which we obtain the credible intervals (CI) of the modulation parameters *K*^mod^ and *ϕ*^mod^. To estimate the posterior distribution, we sample from the posterior distributions given by (i) the state space oscillator model and (ii) the parametric PAC model.

(i) The Kalman Filter used in the *r*^*th*^ E-Step (see Supplementary Materials 9.2.1) of the EM algorithm provides the following moments, for *t*, *t*′ = 1‥*N*:

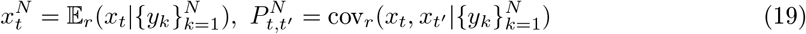

Therefore, we can sample *l*_1_ times series: 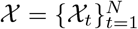 using:

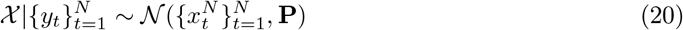

where **P** is a 4*N* × 4*N* matrix whose block entries are given by 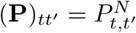.

(ii) For each 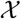, we use equation (11) to compute the resampled slow oscillation’s phase *φ* and fast oscillation’s amplitude 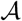. We then use equation (17) to draw *l*_2_ samples from 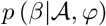. As a result, we produce *l*_1_ × *l*_2_ samples to estimate:

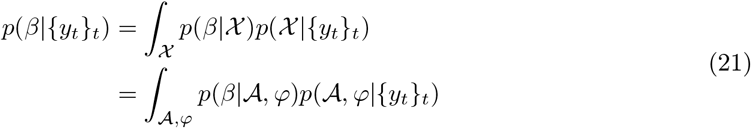

We finally construct CI around *β*^SSP^ using an *L*_2_ norm and in turn derive the CI of *K*^mod^ and *ϕ*^mod^ (Fig.1 h,i).

#### 8.2.4 A Second State-Space Model to Represent Time-Varying PAC

We segment the time series into multiple non-overlapping windows of length *N* to which we apply the previously described analysis. We hence produce 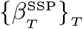, a set of vectors in ℝ^3^ accounting for the modulation where *T* denotes a time window of length *N*.

A second state-space model can be used to represent the modulation dynamics. Here we fit an autoregressive (AR) model of order *p* with observation noise to the modulation vectors 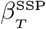 across time windows. It yields the double State Space PAC estimate (dSSP):

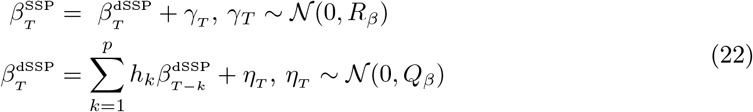

We proceed by solving and optimizing Yule-Walker type equations numerically (see Supplementary 9.4) and we select the order *p* with Bayesian Information Criterion [28]. Finally, we can use the fitted parameters to filter the *l*_1_ × *l*_2_ resampled parameters to construct a CI for 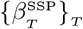 when necessary.

#### 8.2.5 Equivalence

To better compare standard techniques with the SSP, we derive an approximate expression for the PAM under our parametric model (Supplementary Materials 9.5). For a window *T*:

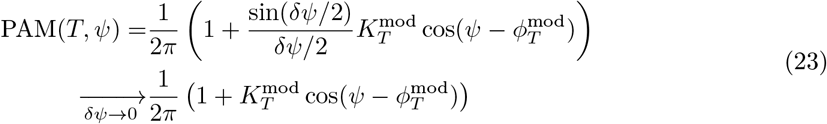

### 8.3 Data Sets

#### 8.3.1 Experimental design and procedure

##### a) Human EEG

We analyzed human EEG data during loss and recovery of consciousness during administration of the anesthetic drug propofol. The experimental design and EEG preprocessing have been extensively described in [23]. Briefly, 10 healthy volunteers (18-36 years old) were infused increasing amounts of propofol spanning 6 target effect site concentrations (0, 1, 2, 3, 4, and 5 *μ*g.L^−1^). Infusion was computer controlled and each concentration was maintained for 14 minutes. To monitor loss and recovery of consciousness behaviorally, subject were presented with an audio stimulus (a click or a verbal command^2^) every 4 seconds and had to respond by pressing a button. The probability of response and associated 95% credible intervals were estimated using Monte-Carlo methods [46] to fit a state space model to these data. Finally, EEG data were pre-processed using an anti-aliasing filter and downsampled to 250Hz.

We computed spectrograms of the EEG using the parametric expression associated with oscillation decomposition (derived in Supplementary Materials 9.1). Standard techniques for PAC analysis were applied on 6 and 120 second windows for which alpha and slow component were assumed to be known and extracted using bandpass filters around 0.1-1Hz and 9-12Hz. Significance for the standard PAC method was assessed using 200 random permutations.

##### b) Rat LFP

Rat LFP dataset was generously shared by Tort et al. [4]. Data were recorded from the CA3 region of the dorsal hippocampus of rats as they learned a spatial recognition task. Signal was sampled at 1000Hz, bandpassed from 1 to 300Hz and binned into non overlapping 2 second time windows. The standard PAC analysis was performed using 6-10Hz and 30-55Hz filters to extract theta and gamma components, respectively. To replicate the original results, modulation indices were averaged over 20 trials.

#### 8.3.2 Simulations

We tested our algorithm on simulated datasets generated by different models. We constructed each simulated signal by combining unit variance Gaussian noise, a slow oscillation centered at *f*_s_ (=1Hz unless stated otherwise), and a modulated fast oscillation centered at *f*_f_ (=10Hz unless stated otherwise). It is important to note that we chose to generate these simulated signals using a method or “model class” that was different from the state space oscillator model we use to analyze the data. For standard processing, significance was assessed with 200 random permutations, *f*_s_ and *f*_f_ were assumed to be known, and components were extracted with bandpass filters with pass bands set to 0.1-1Hz for the slow component and 8-12Hz for the fast component.

##### a) Simulating the Slow Oscillation

Neural oscillations are not perfect sinusoids and instead have a broad band character. Using the approach described in [24], we simulated a broad band slow oscillation by convolving (filtering) independent identically distributed Gaussian noise with the following impulse response function:

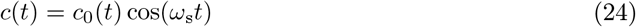

where *ω*_s_ = 2*πf*_s_, *c*_0_ is a Blackman window of order 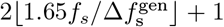. The smaller the slow frequency bandwidth 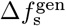, the closer the signal is to a sinusoid. When necessary, we additionally use a *π*/2 phase-shifted filter: 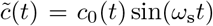 to model an analytic slow oscillation 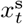 from which we deduce the phase 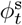. The resulting series is finally normalized such that its standard deviation is set to *σ*_s_.

##### b) Simulating the Modulation

To assess the temporal resolution of our method and the standard method, we generated simulated data sets with different rates of time-varying modulation. First, to construct the modulated fast oscillation, we constructed a fast oscillation centered at *ω*_f_ = 2*πf*_f_ and normalized to *σ*_f_ as described above and modulated it by:

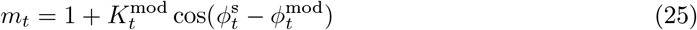

Here, 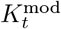 and phase 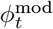 are time varying and follow the dynamics illustrated in Fig. 5 (Section 2.4) and in Fig.11 (Supplementary Materials 9.6.3). Representative simulated EEG signal traces for different generative parameters are illustrated in Supplementary Fig. 12.

We also generated simulated data using an alternative modulation function (Fig. 6 and Supplementary Fig. 13, 14 and 15)

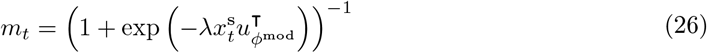

where 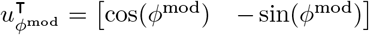, described previously in described in [24].

##### c) Simulated Signals with Abrupt Changes

Signals with abrupt or sharp transitions can lead to artefactual phase-amplitude modulation [19]. To assess the robustness of our state-space PAC method under such conditions, we used a Van Der Pol oscillator to generate a signal with abrupt changes. Here, the oscillation *x* is governed by the differential equation:

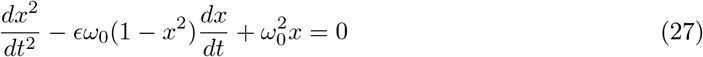

Equation (27) was solved using Euler method with fixed time steps.

## 9 Supplementary Materials

### 9.1 Power Spectral Density for the State Space Oscillator Process

In this section, we derive the parametric expression for the power spectral density (PSD) of an oscillation 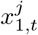 by building an autoregressive moving average process (ARMA) with the same spectral content. For convenience, we will drop the index *j* in what follows. First, we note that an oscillation is asymptotically second order stationary. Let us compute its autocovariance sequence. Since 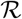 is a rotation matrix 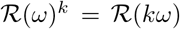 and 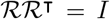. Therefore, from equation (8), it comes:

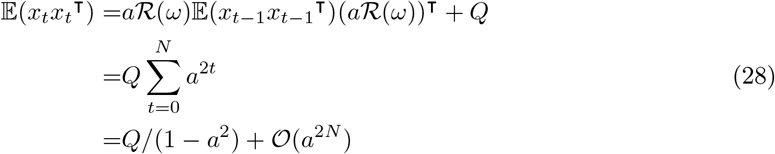

and:

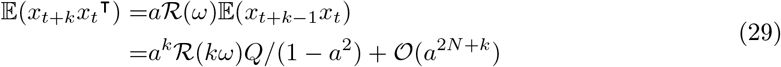

We can hence write, for *t* = 1‥*N*, *k* = 0‥*N* − *t*, 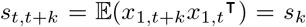. As a consequence, an oscillation can be approximated by a second order stationary process, and in virtue of the Wiener-Khinchin theorem [36], its theoretical power spectral density is:

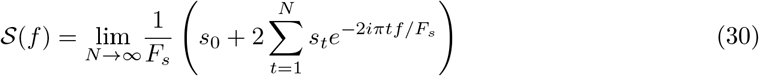

We now consider the ARMA(2,1):

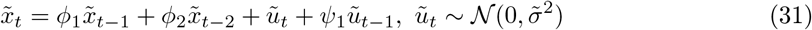

to which we impose, for *t* = 1‥*N*, *k* = 0‥*N* − *t*: 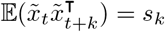. It follows that:

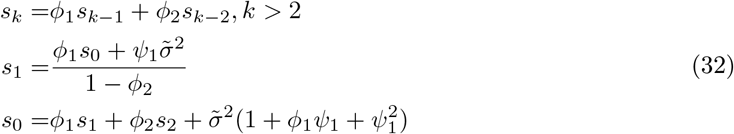

Taking:

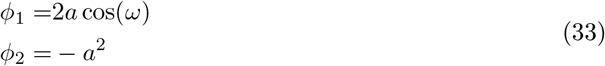

satisfies the first equality of equation (32). The remaining conditions can then be rewritten:

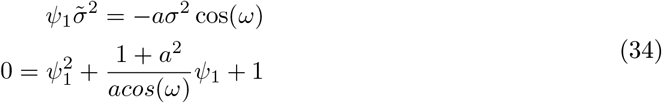

from which we deduce 2 two negative roots 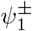 ultimately leading to the same autocovariance series. Overall, we choose:

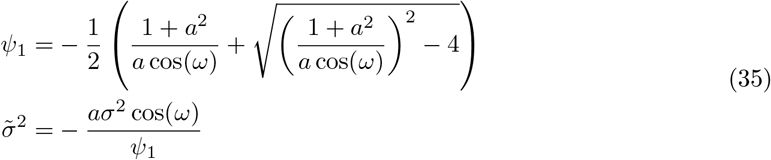

Applying the discrete Fourier transform to equation (31) yields:

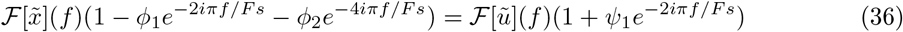

since 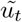 is Gaussian noise, 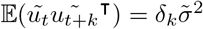. Therefore, our ARMA(2,1) PSD is:

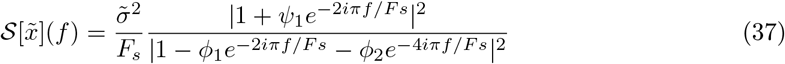

Finally, the PSD of an oscillation centered in *f*_0_ is;

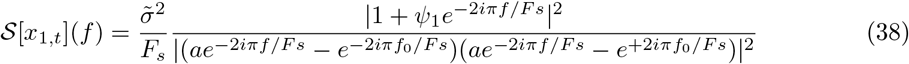

### 9.2 An Expectation-Maximization (EM) Algorithm for Independent and Harmonic Oscillation Decompositions

Since all noise terms are assumed to be additive Gaussian, the complete data log likelihood for one time window of length *N* is:

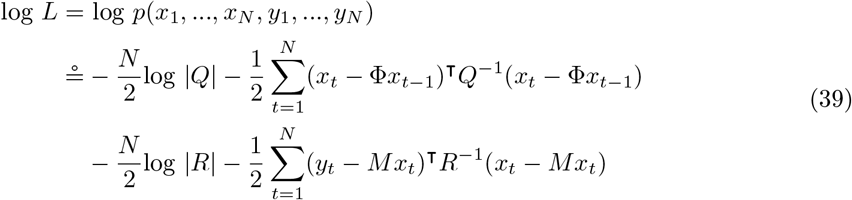

We wish to maximize log *L* with respect to Θ = (Φ, *Q*, *R*) but we do not have access to the hidden oscillations *x*_*t*_. We use an expectation maximization algorithm to alternatively and iteratively estimate (E-Step) and maximize (M-Step) the log likelihood. At iteration *r*, we use the Kalman filter to estimate *x*_*t*_ given a set Θ_*r*_ which gives us access to:

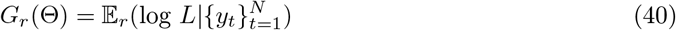

Then, we deduce Θ_*r*+1_:

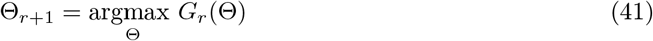

#### 9.2.1 Kalman Filter and Fixed Interval Smoother Estimates

We use the Kalman filter to estimate the hidden oscillations given the observations and the model parameters. They first predict the state at the next time point, then compare that prediction to the observation, and finally produce an updated estimate based on the most recently observed data. Given the full observation time series, we can apply backward smoothing to refine the update to account for the full observation series (i.e., fixed interval smoothing).

For *t*, *t*_1_, *t*_2_ = 1‥*N*, we note:

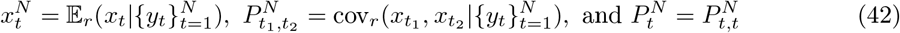

and compute those quantities using the forward smoothing algorithm:

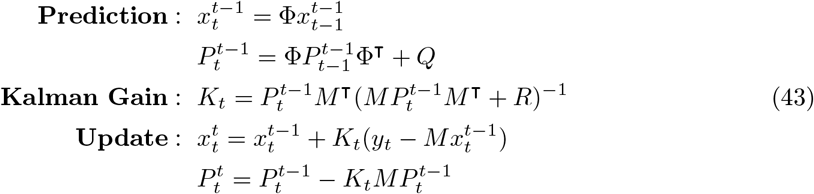

and a set of backward recursions [48]. For *t* = *N*‥1, and *t*_1_ < *t*_2_:

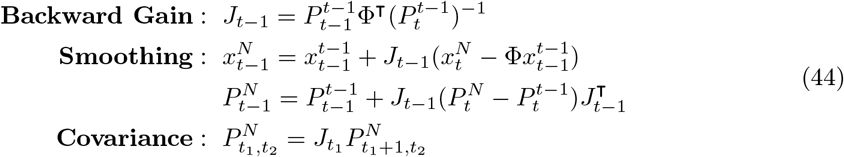

#### 9.2.2 E-Step

Here we describe *G*_*r*_(Θ) for the state space oscillation decomposition with harmonic components. An single oscillation is defined by a rotation matrix 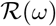, a scaling parameter *a* and a process noise covariance matrix *Q* = *σ*^2^*I*_2×2_. In what follows, we consider *d* independent oscillations, which are respectively associated to *h*_1_,…, *h*_*d*_ harmonics. For *j* = 1‥*d*, an oscillation with fundamental frequency *ω*_*j*_ is the sum of *h* = 1…*h*_*j*_, harmonics respectively defined by 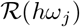, *a*_*j*,*h*_ and 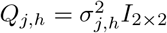. We note 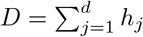 the total number of oscillatory components.

For *V* ∈ ℝ^2*D*×2*D*^, *j* = 1‥*d* and *h* = 1‥*h*_*j*_, we note *V*_*j*,*h*_ the 2 by 2 diagonal block associated with the 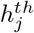 harmonic of oscillation *j*. Φ and *Q* are block diagonal matrices whose diagonal blocks are 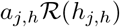 and *Q*_*j*,*h*_:

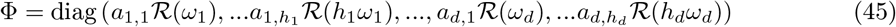

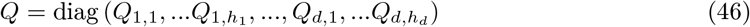

Additionally, we will use *M* = [1 0 1 0 … 0] ∈ ℝ^2*D*^ and for *U* ∈ ℝ^2×2^, we note: rt(*U*): = *U*_21_ − *U*_12_ and tr(*U*) = *U*_11_ + *U*_22_.

Taking the conditional expectation of the log likelihood log *L* at iteration *r* for a fixed set of parameter Θ_*r*_ = (Φ, *Q*, *R*)_*r*_, we obtain [21]:

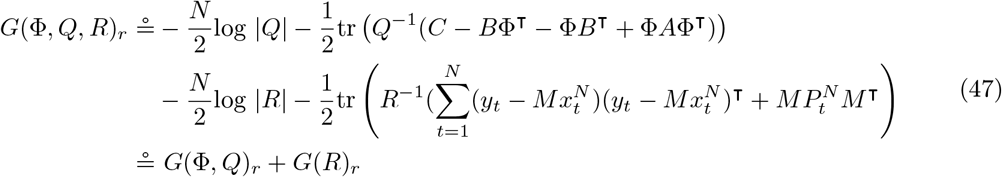

where:

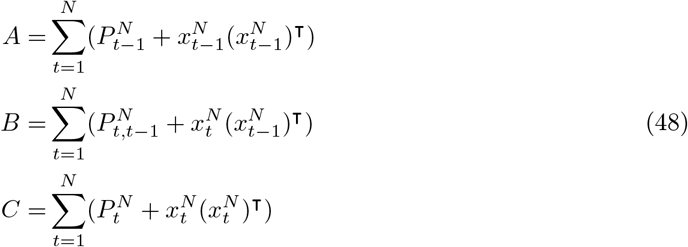

#### 9.2.3 M-Step

We can maximize *G*_*r*_ with respect to *R* and (Φ, *Q*) independently. We have:

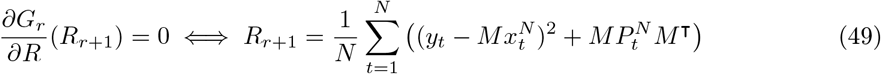

Since *Q* is (block) diagonal, we can write:

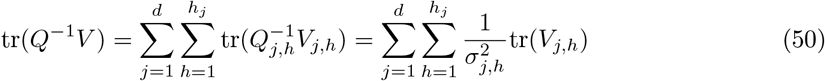

*A* is symmetric and Φ is a block diagonal matrix whose element are 2 × 2 rotation matrices, we develop equation (47) and obtain:

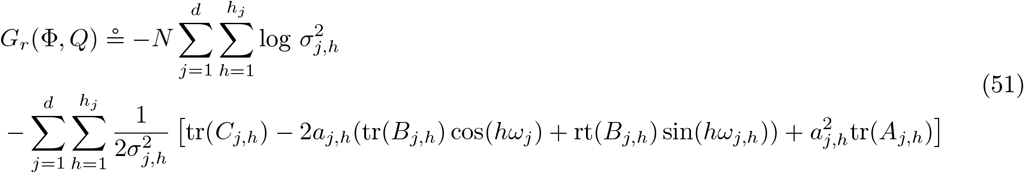

Differentiating with respect to process noises covariances 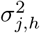, scaling parameter *a*_*j*,*h*_ and fundamental frequencies *ω*_*j*_ yields:

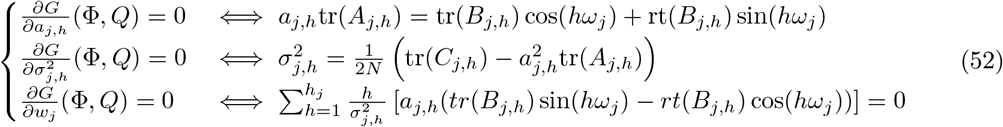

We inject the upper equations of (52) into the third one and we note:

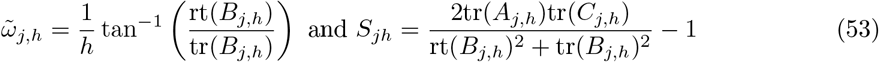

Using trigonometric identities, equations (52) can finally be rewritten:

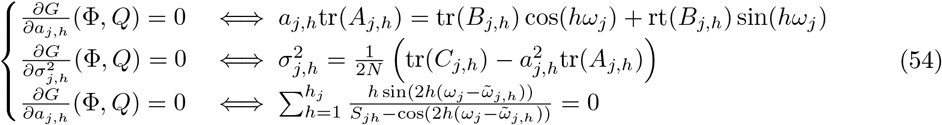

Overall, for *j* = 1‥*d*, if *h*_*j*_ > 1, we numerically solve for *ω*_*j*_ using equation (54) and deduce *a*_*j*,*h*_ and 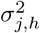 for *h* = 1‥*h*_*j*_.

If *h*_*j*_ = 1, we immediately have:

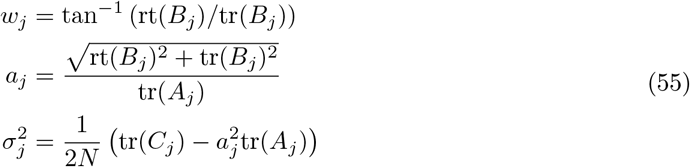

### 9.3 Linear Regression: priors, hyperparameters, and normalizing constants

As in [45], we use 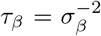 and we assume that the likelihood of the model defined in equation (14) is:

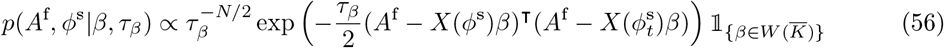

where 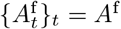 and 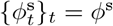. The conjugate prior is:

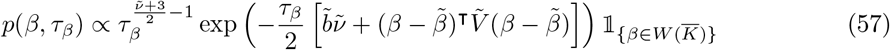

We choose prior hyperparameters 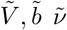 and 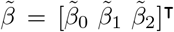 to convey as little information as possible on the phase and the strength of the modulation. Marginalizing equation (57) over *τ*_*β*_, yields a truncated multivariate t-distribution:

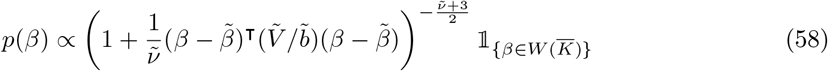

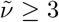 insures that the multivatiate-t variance is defined. It is: 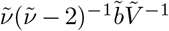. Then, we consider the independent random variables 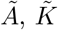 and 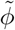 such that:

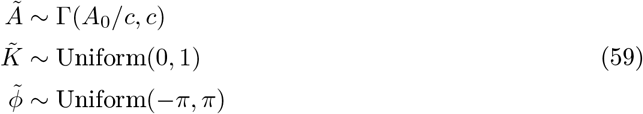

We note 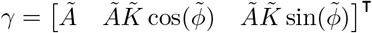, use 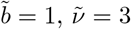 and we define 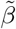, and 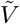 such that:

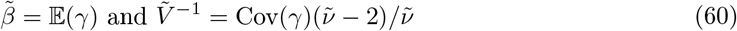

Additionally, we notice that if 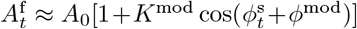 and since 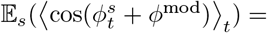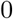 (where 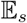 represents an average over trials or windows and 〈.〉_*t*_ is a temporal average across a given window), 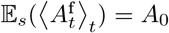 and 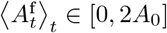. Therefore, it is reasonable to use 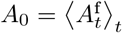 and *c* = 1. Overall:

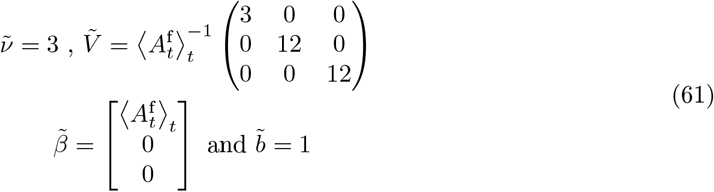

Posterior parameters are then given by:

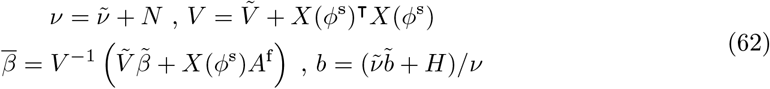

Where:

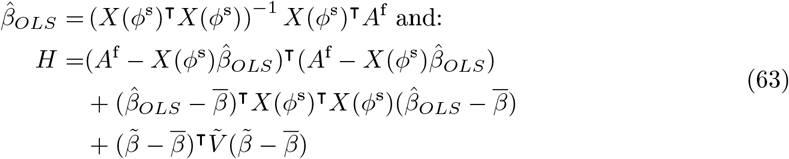

We deduce the posterior distribution:

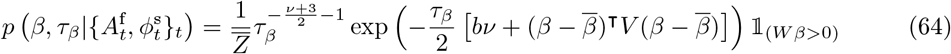

The normalizing constant 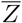 is obtained by integration and is:

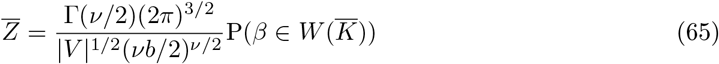

where 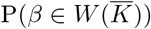 is computed using the multivariate t-distribution of parameters *v*, 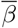 and *b*^−1^*V*.

Finally, we deduce equation (17) by marginilizing equation (64) over *τ*_*β*_ and have:

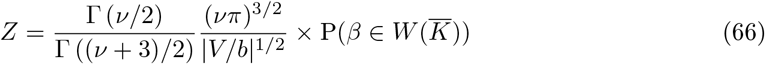

Note that for large samples it might be useful to use:

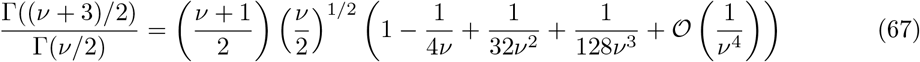

### 9.4 Second State-Space

In this section, for a given AR order *p*, we estimate the parameters *R*_*β*_, *Q*_*β*_ and 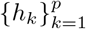 defined equation (22). Let 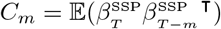 be the autocovariance sequence of the modulation vectors estimated with equation (18). We have:

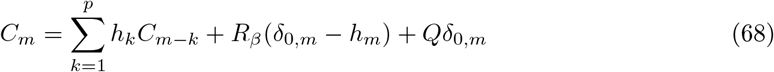

where *δ*_*i*,*j*_ is the the Kronecker delta. Equation (68) can be rewritten:

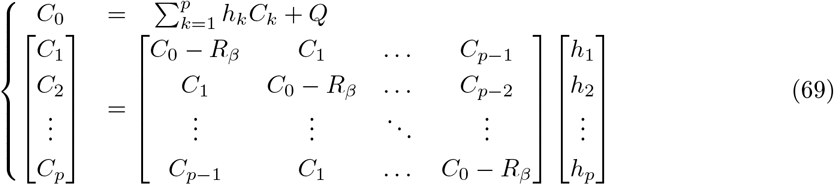

For an observation noise candidate 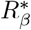, if we can invert equation (69), we immediately access 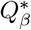 and 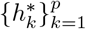. Using the Kalman Filter, we hence deduce the likelihood of the candidate model as in [20].

Therefore, we note 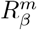 the smallest eigenvalue of the Toeplitz matrix **C** = (*C*_|*i*−*j*|_)_*i*,*j*=0‥*p*_, and, numerically ^3^ optimize the model likelihood with respect to 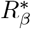 in 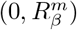, where we know that 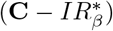 is full rank.

From *R*_*β*_ we get *Q*_*β*_ and 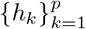 then, once again, we use the Kalman Filter to estimate 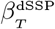.

### 9.5 Modulogram Equivalence

We derive an approximated parametric modulogram for a window of length *δt* = *N*/*F*_*s*_ centered in *τ*. We will use:

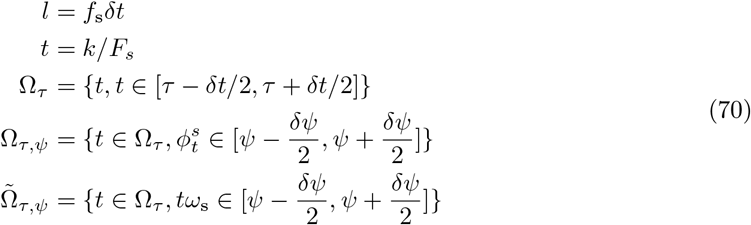

For clarity we will use *t* or *k* without distinction and we remind that:

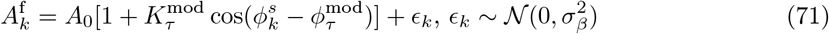

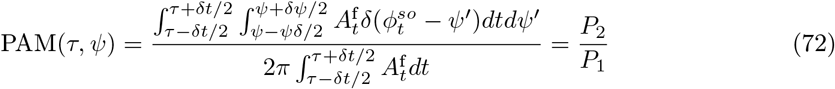

Additionally, we assume that:
- for *k* ∈ Ω_*τ*_, 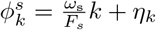, where 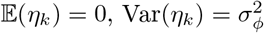 and 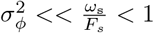
- and, for simplicity, for *ψ* ∈ [−*π*, *π*], for all *h*: ℝ^+^ → ℝ smooth, 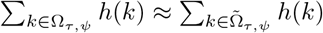

From the central limit theorem, 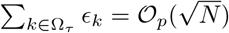 and 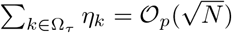.

We hence have:

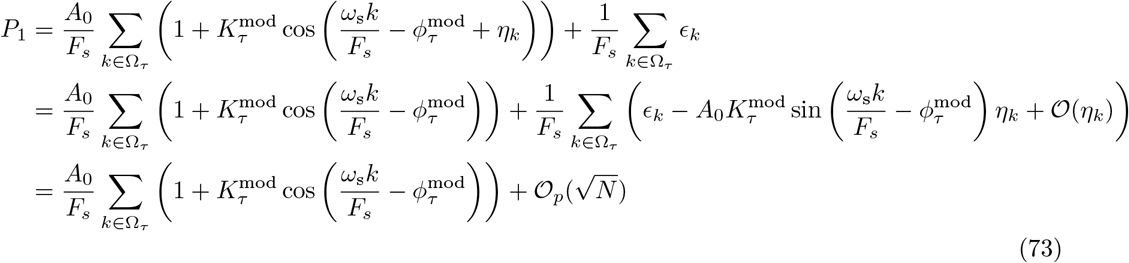

But 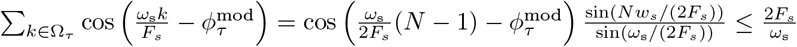. Therefore:

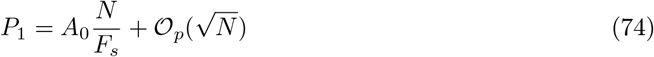

On the other hand:

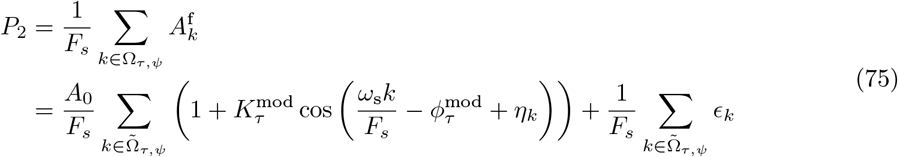

But 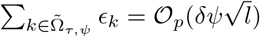 and *l* ∝ *N* so:

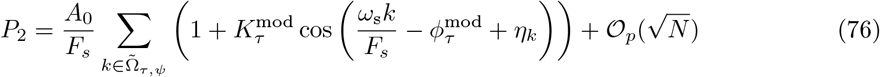

Using the same argument as the one detailed above we get:

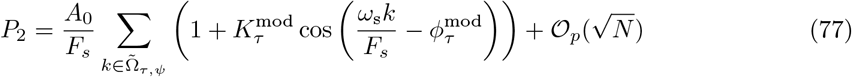

Additionally:

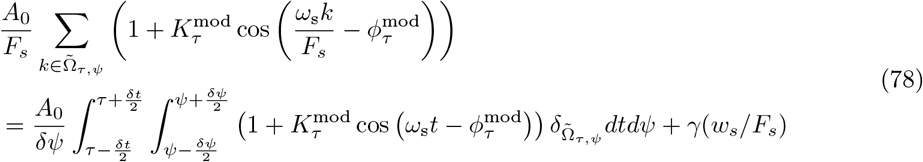

Where *γ* is a function such that 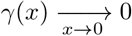. Since:

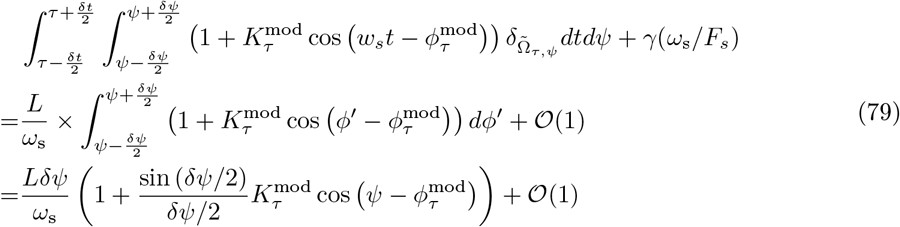

For 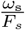 “sufficiently small”, we can write:

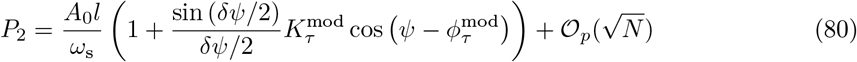

Finally:

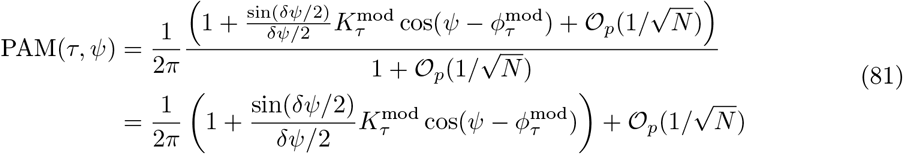

### 9.6 Initialization of the EM algorithm

Although EM insures convergence, the log likelihood which is to be maximized is not always concave [20]. To address this issue, Matsuda and Komaki initialize a signal composed of *d* oscillations with the parameters of the best autoregressive (AR) process of order *p* ∈ [|*d*, 2*d*|]. Nevertheless, because of the electrophysiological signal’s aperiodic component, such procedure might bias the initialization. Indeed, the aperiodic components are usually described by a 1/*f*^*χ*^ power-law function [43] [44] which might be regressed by the AR process. In such cases, the initialization could fail to account for an actual underlying oscillation.

To help mitigate this potential problem, we adapt Haller, Donoghue and Peterson’s FOOOF algorithm [25] to the state space oscillation framework. Our initialization algorithm aims to disentangle the oscillatory components from the aperiodic one before fitting the resulting spectra^4^ with the parametric PSD of the oscillation (equation (38)).

The power spectral density (PSD) for the observed data signal *y*_*t*_ is estimated using the multitaper method [36]. We set the frequency resolution *r*_*f*_ (typically to 1Hz) which yields the time bandwidth product 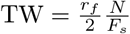. The number of taper *K* is then chosen such that *K* ≪ ⌊2TW⌋ − 1.

First of all, we estimate the observation noise *R*_0_ (used to initialized *R*) using:

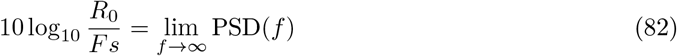

and we remove this offset from the PSD.

#### 9.6.1 Regressing out the non oscillatory component

The aperiodic signal PSD in dB, at frequencies *f* is then modeled by:

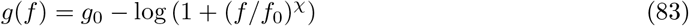

*χ* controls the slope of the aperiodic signal, *g*_0_ the offset and *f*_0_ the “knee” frequency. A first pass fit is applied to identify the frequencies corresponding to non oscillatory components: only *f*_0_ is fitted while *χ* and *g*_0_ are respectively set to *χ* = 2 and *g*_0_ = PSD(*f* = 0). (Fig. 9-a). We fix a threshold (typically 0.8 quantile of the residual) to identify frequencies associated to the aperiodic signals (Fig. 9-b).

**Figure 9:**
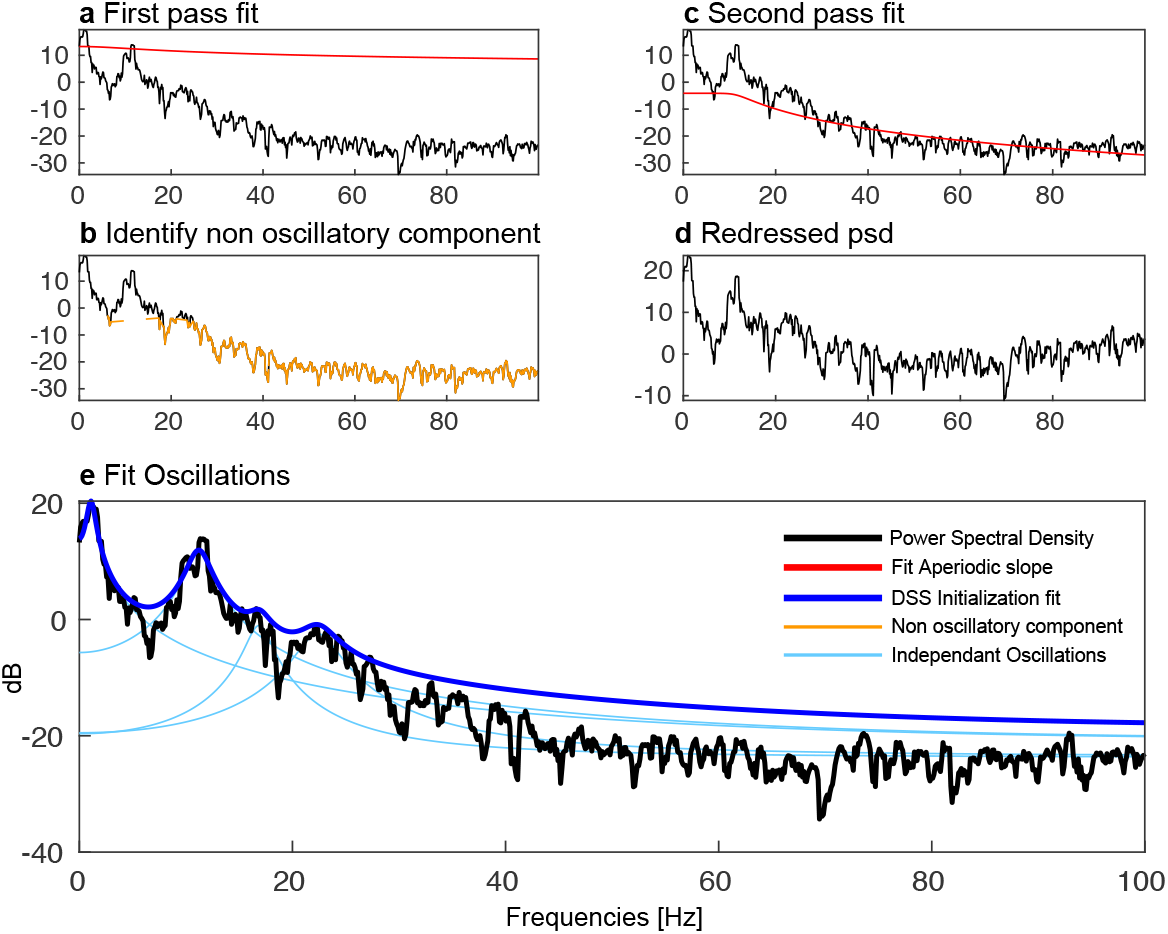
Steps for the initialization procedure. A first pass fit is applied to the raw multitaper power spectral density (PSD) estimate (a). We remove this fit from the raw PSD and fix a threshold to identify non-oscillatory components (b). A second pass fit is applied (c) which yields a redressed PSD (d). We then fit the p arametric e xpression of the PSD for a fixed number of oscillations (d). The fitted parameters are then used to initialize the EM algorithm.

A second pass fit is then applied only on those frequencies from which we deduce *g*_0_, *f* _0_ and *χ* (Fig. 9-c). We remove *g*(*f*) from the raw PSD in dB and use it for the second step of the algorithm (9-d).

#### 9.6.2 Oscillation Initialization

From the redressed PSD, we fit a given number *d*_0_ (e.g *d*_0_ = 4) of independent oscillations using the theoretical PSD given in equation (38). To do so, we identify PSD peaks of sufficient width (wider than *r*_*f*_/2) before fitting an oscillation theoretical spectra in an eighborhood of width 2*r*_*f*_ around this peak. For oscillation *j*, we deduce (*f*_*j*_)_0_, (*a*_*j*_)_0_ and 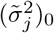. Since 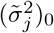 represents the offset of a given oscillation after removing the aperiodic component, we adjust it to estimate 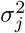:

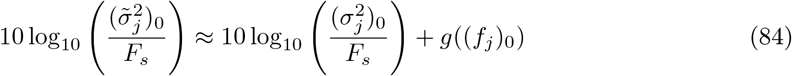

The resulting spectra PSD_*j*_ are then subtracted and the process is repeated until all oscillations are estimated (9-e, blue). We finally estimate the power P_*j*_ of an an oscillation *j* in the neighborhood of (*f*_*j*_)_0_ and estimate its contribution to the total power P_0_ by 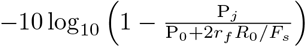.

Oscillation are sorted and the resulting parameters are used to initialize the EM algorithm with the *d* ∈ [|1, *d*_0_|] first oscillations.

#### 9.6.3 Additional Results

In this section we present additional results to support the validation and comparison of our algorithm:

Fig. 10 is the PAC profile of another subject infused with increasing target effect site concentration of propofol.

**Figure 10:**
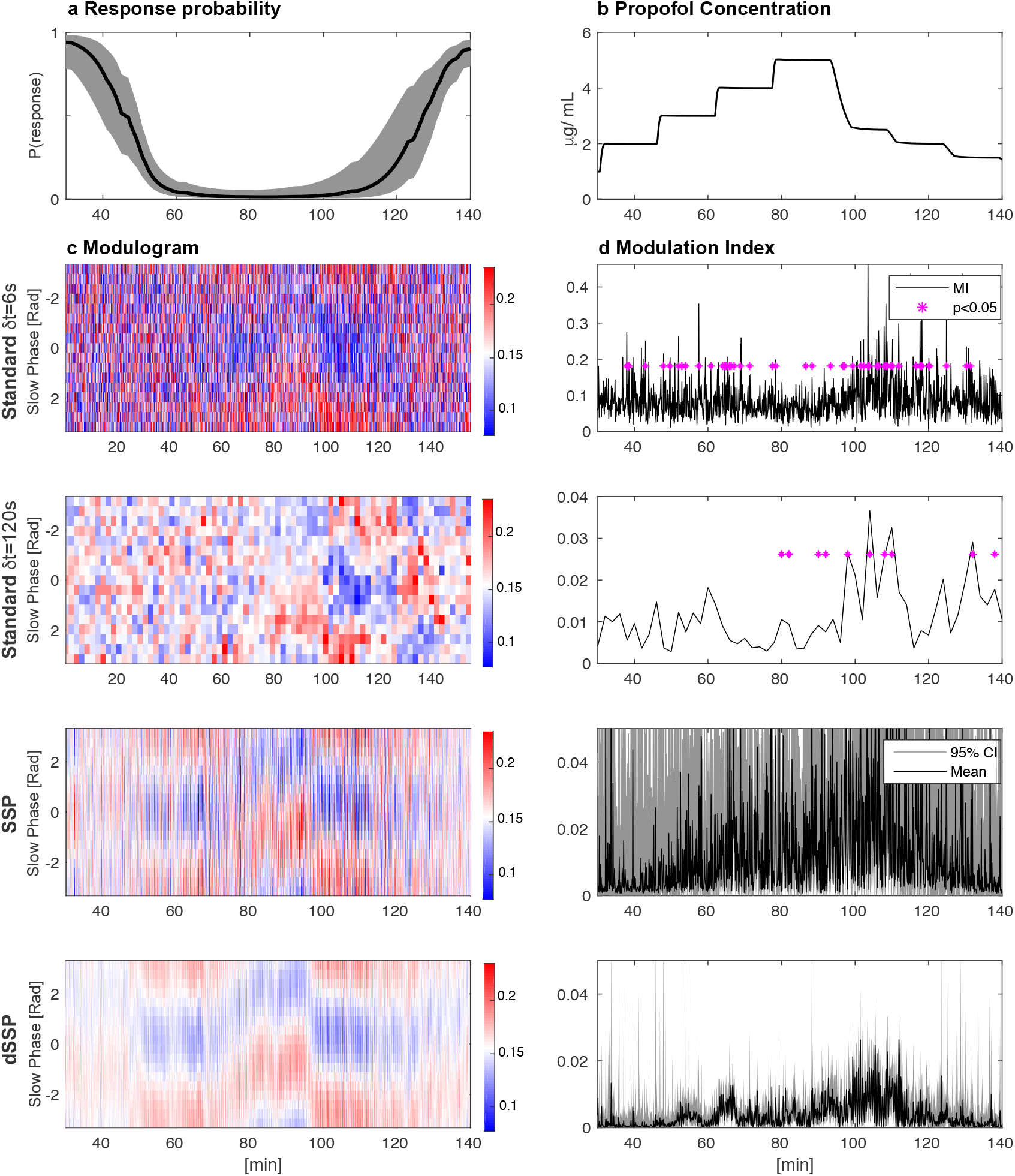
Phase Amplitude Coupling profile of another subject infused with increasing target effect site concentration of propofol. Left: response probability curves (a) aligned with modulograms (c) (distribution of alpha amplitude with respect to slow phase) computed with standard (top) and parametric (bottom) techniques. Right: propofol infusion target concentration (b) aligned with corresponding Modulation Indices (d). Standard technique significance was assessed using 200 random permutations and CI where estimated using 200 × 200 samples

Fig.11 is a comparison of the modulation dynamic estimation between standard analysis and our dSSP.

**Figure 11:**
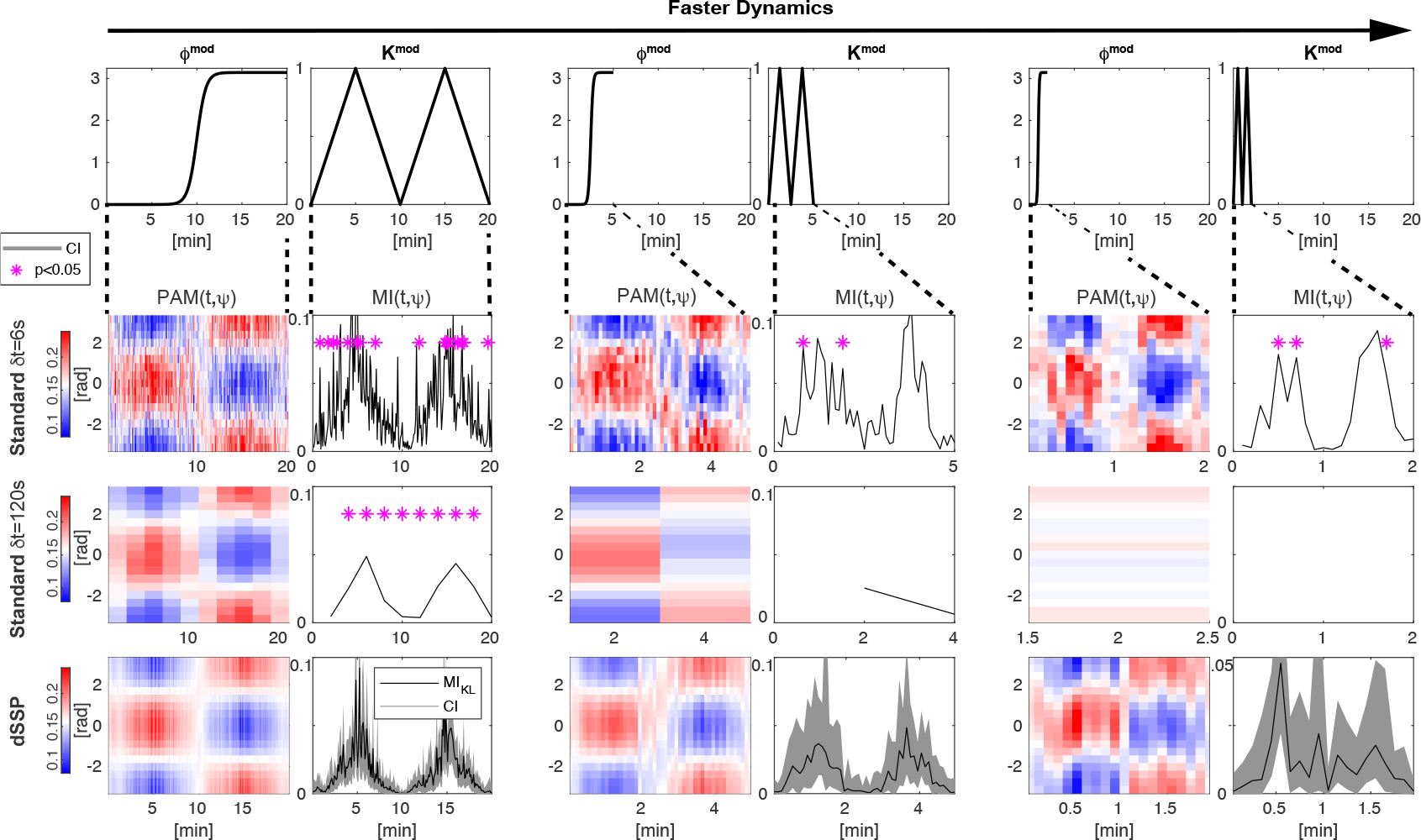
Comparison of the modulation estimates using standard methods and our new dSSP method. Slow and fast oscillations were generated by filtering white noise around *f*_s_ = 1Hz and *f*_s_ = 10Hz with 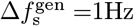 and normalized to standard deviation *σ*_s_ = 1 and *σ*_f_ = 0.8. The time scale over which *K*^mod^ and *ϕ*^mod^ changed varied between 20 minutes to 5 and 2 minutes. See Fig. 12 for typical signal traces.

Fig.12 are typical 6s signal traces of signal generated with equation (25).

**Figure 12:**
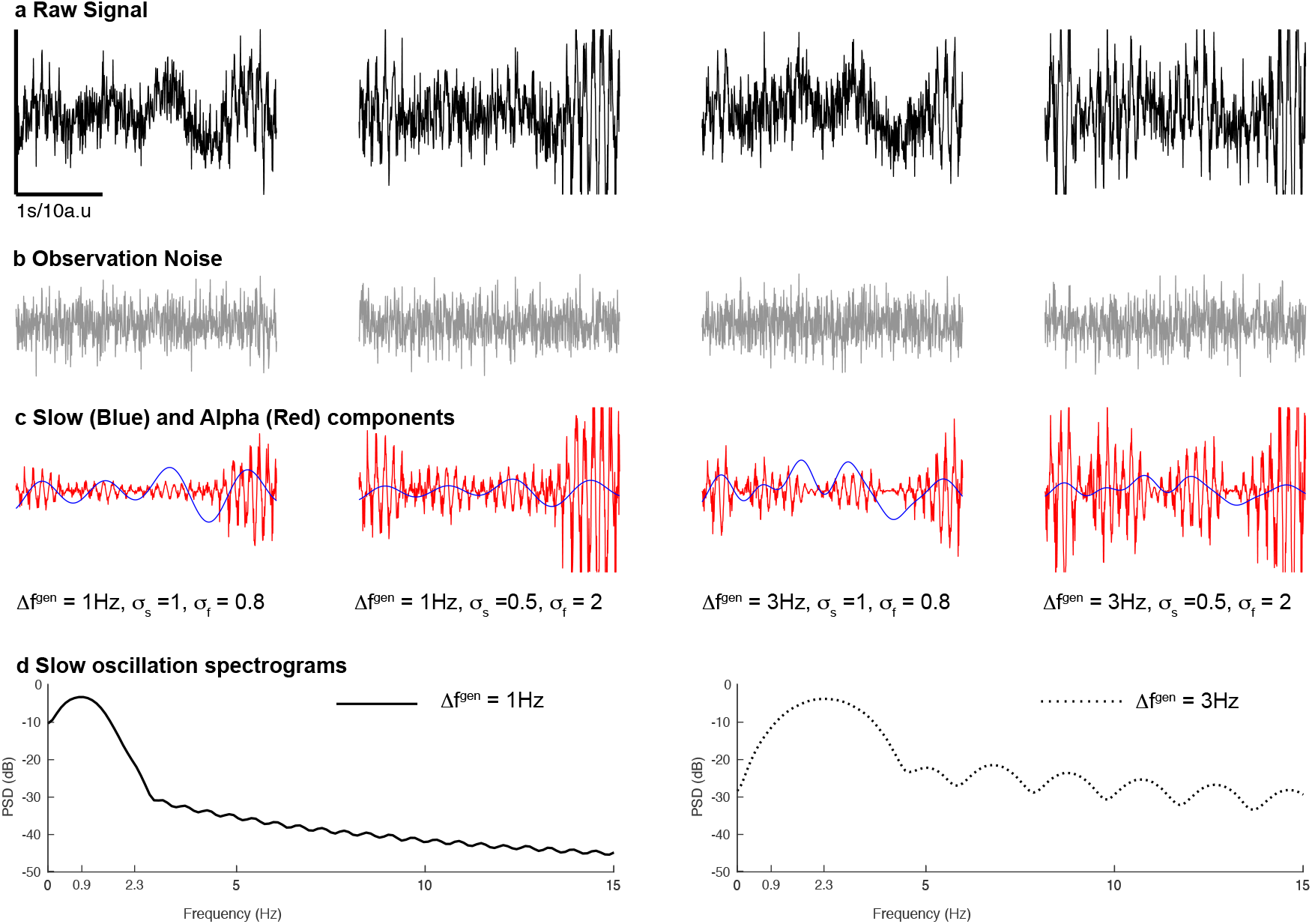
Typical 6s signal traces generated with equation (25) for 4 different conditions. The raw signal (a) is the sum of the observation noise (b), slow, and alpha oscillations (c). We show slow frequency multitaper PSD (TW=4, K=5 tapers) for different values of 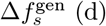

Fig.13, 14 and 15 Are the modulation phase (*ϕ*^mod^) and frequency (*f*_s_, *f*_f_) recovery estimation and comparison between standard methods, DAR and SSP.

**Figure 13:**
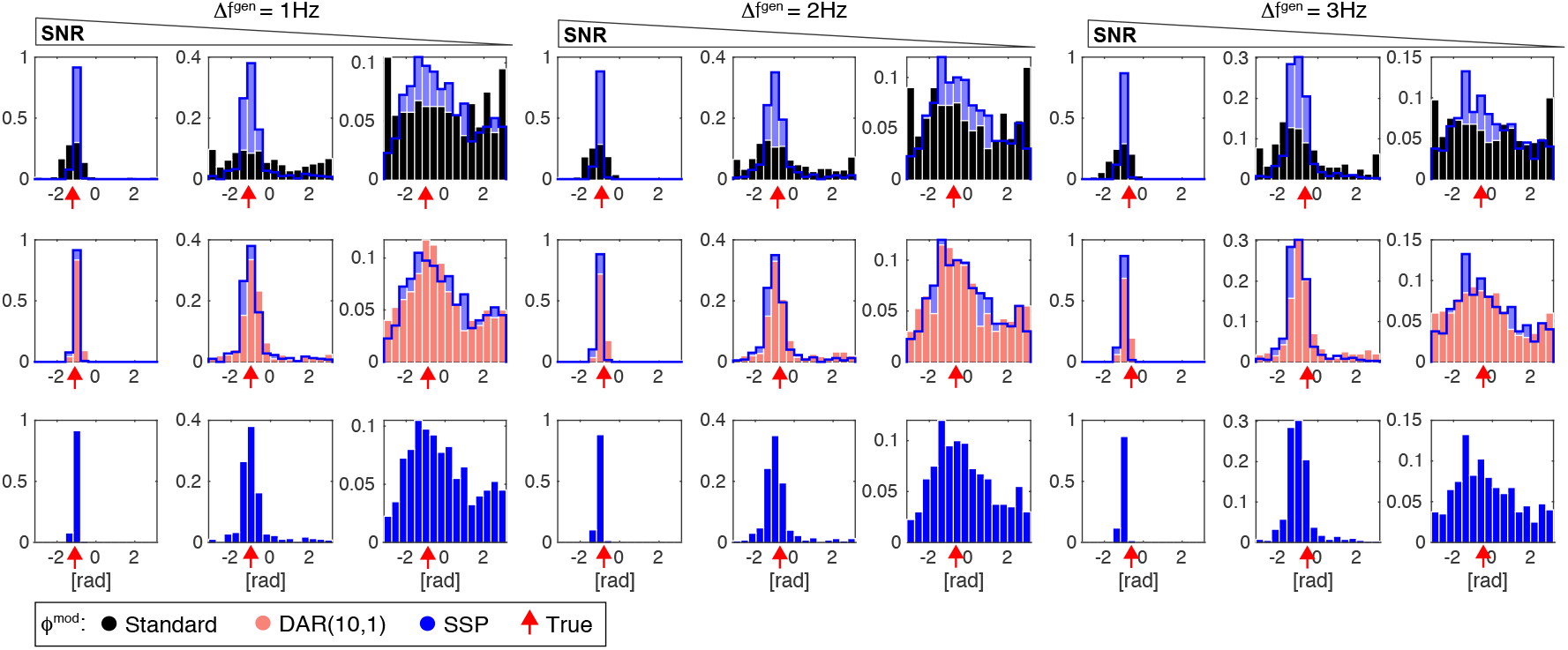
Modulation phase *ϕ*^mod^ estimation and comparison between standard methods (black), DAR (pink) and SSP (blue). 400 windows of 2 seconds were generated with: a slow oscillation (filtered from white noise around *f*_s_ = 3Hz with 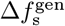 and normalized to standard deviation *σ*_s_) and a modulated fast oscillation (*ϕ*^mod^ = −*π*/3, modeled with a sinusoid *f*_s_ = 50Hz and normalized to *σ*_f_). We added unit normalized Gaussian noise and we used 3 Signal To Noise Ratio (SNR) conditions ((*σ*_s_, *σ*_s_) = (2, 1.5), (1, 0.6) and (0.7, 0.3)). We show typical signal traces for these different conditions in Supplementary Materials Fig 15.

**Figure 14:**
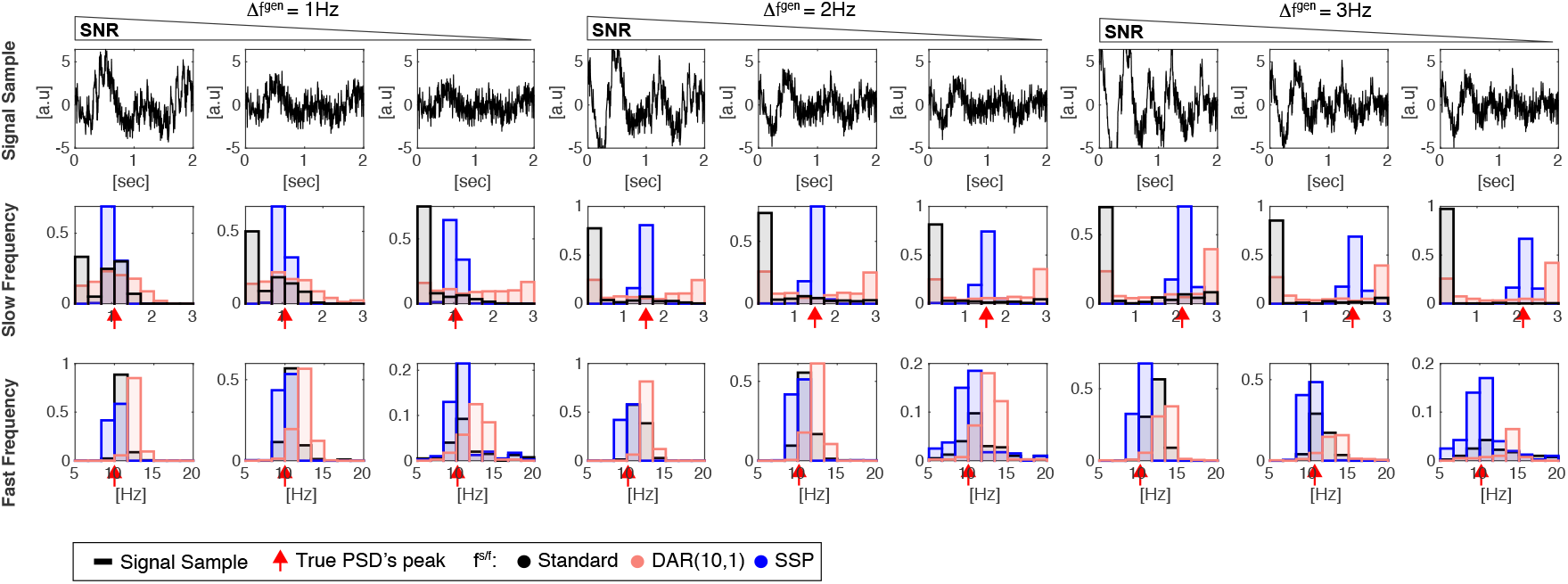
Slow and Fast oscillation recovery using different algorithms: standard methods (black), DAR (pink) and SSP (blue). 400 windows of 6 seconds were generated with: a slow oscillation (filtered from white noise around *f*_s_ = 1Hz with bandwidth 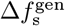 and a modulated fast oscillation (*ϕ*^mod^ = −*π*/3, modeled with a sinusoid *f*_s_ = 10Hz and normalized to *σ*_f_). We added unit normalized Gaussian noise and we used 3 Signal To Noise Ratio (SNR) conditions ((*σ*_s_, *σ*_s_) = (2, 1.5), (1, 0.6) and (0.7, 0.3)). We report typical signal traces for the different conditions (top), slow oscillation recovery alongside the true slow frequency PSD (middle), and fast frequency recovery (bottom). The red arrow indicates the true multitaper PSD (TW=4, K=5 tapers) peak for each oscillation.

**Figure 15:**
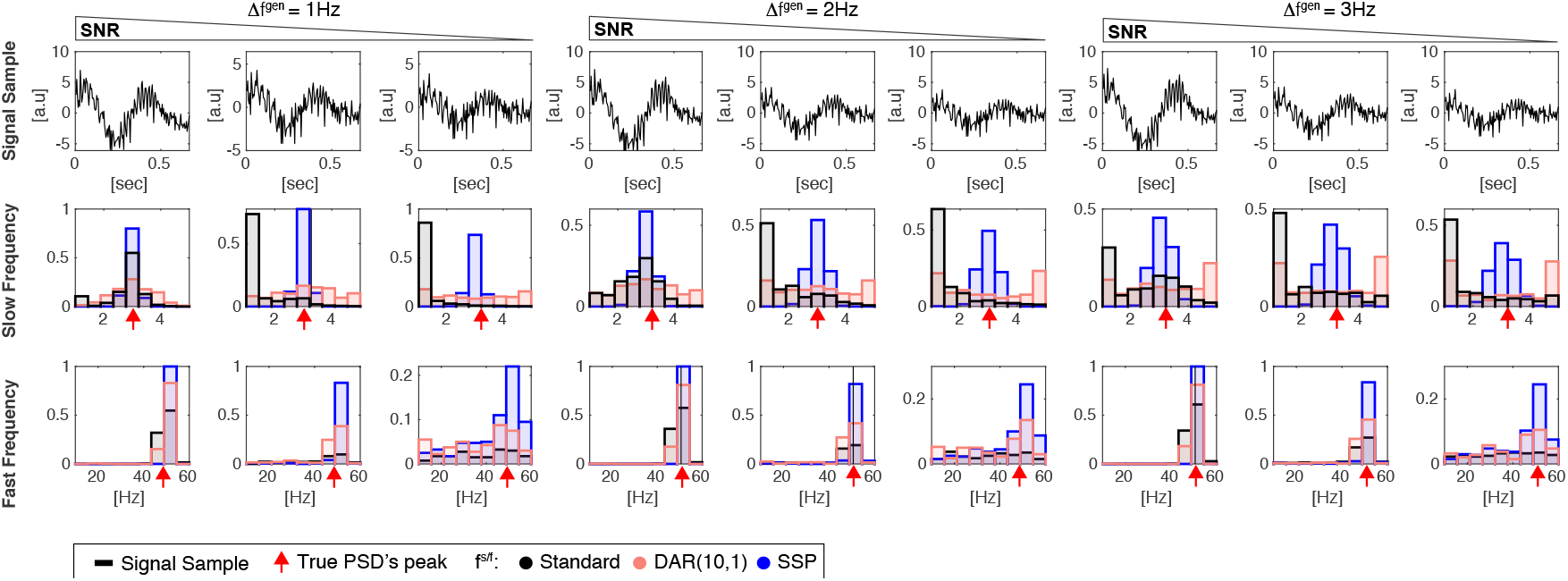
Slow and Fast oscillation recovery using different algorithm: standard methods (black), DAR (pink) and SSP (blue). 400 windows of 2 seconds were generated with: a slow oscillation (filtered from white noise around *f*_s_ = 3Hz with bandwidth 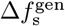 and normalized to standard deviation *σ*_s_) and a modulated fast oscillation (*ϕ*^mod^ = −*π*/3, modeled with a sinusoid *f*_s_ = 50Hz and normalized to *σ*_f_). We added unit normalized Gaussian noise and we used 3 Signal To Noise Ratio (SNR) conditions ((*σ*_s_, *σ*_s_) = (2, 1.5), (1, 0.6) and (0.7, 0.3)). We report typical signal traces for the different conditions (top), slow oscillation recovery alongside the true slow frequency PSD (middle), and fast frequency recovery (bottom). The red arrow indicates the true multitaper PSD (TW=4, K=5 tapers) peak for each oscillation.

## Footnotes

1 unless stated otherwise *Fs* = 250Hz and *N/Fs* = 6s

2 here, we only report the response to the verbal command

3 golden section search and parabolic interpolation

4 All fits in this initialization procedure use interior point methods to minimize *L*_2_ norms.

